# A genome-wide analysis identifies DIDO3 binding at active chromatin and topological boundaries in mouse stem cells

**DOI:** 10.1101/2021.09.27.462041

**Authors:** Tirso Pons, François Serra, Agnes Fütterer, Karel H. M. van Wely, Florencio Pazos, Cristina Pacios, Ana Cayuela López, Carlos Oscar S. Sorzano, Alfonso Valencia, Carlos Martínez-A

## Abstract

While chromatin structure influences gene expression, how chromatin readers exert transcriptional control is still not fully understood. DIDO3, an isoform of the Death Inducer Obliterator protein, has been implicated in cell differentiation and cancer, yet its precise genomic functions are unclear. Here we integrated ChIP-seq, RNA-seq and Hi-C analyses in mouse embryonic stem cells to reveal that DIDO3 predominantly binds to RNA polymerase II at topological domain boundaries and chromatin loops. We demonstrate that DIDO3 interacts with the architectural protein CTCF and that the N-terminal region of DIDO3 (DIDO3_1-528_) also co-localizes with the histone marks H3K4me3 and H3K36me3. DIDO3 influences the expression of genes encoding chromatin remodeling components, as well as those involved in transcriptional dynamics and three-dimensional chromatin organization. These findings suggest that DIDO3 is associated with both transcriptional regulation and chromatin organization, with potential implications in terms of cell differentiation and tumorigenesis.

## INTRODUCTION

The Death Inducer Obliterator gene (*Dido1*) is expressed ubiquitously in all tissues (García-Domingo et al., 1999) and through alternative splicing, it encodes three protein isoforms: DIDO1, DIDO2, and DIDO3 (Fütterer et al., 2005). All three isoforms share a plant homeodomain (PHD) that recognizes the amino-terminal tail of histone H3 (Gatchalian et al., 2013; Rojas et al., 2005), while DIDO2 and DIDO3 also contain a transcription elongation factor S II subunit M (TFIIS_M) domain that mediates binding to RNA polymerase II (RNAPII: Fütterer et al., 2017; Rojas et al., 2005) and a phosphoserine binding (SPOC) domain that acts as a reader of the C-terminal domain of RNAPII (see Figure 1: Appel et al., 2022; Sánchez-Pulido et al., 2004). The combination of these domains suggests that DIDO3 functions both as an auxiliary protein of RNAPII and as a chromatin reader (Tencer et al., 2017). However, how these domains interact and contribute to the regulation of global genomic processes is not yet fully understood (García-López et al., 2024; Fütterer et al., 2024, 2021, 2017, 2012; Gutiérrez et al., 2022; Mora Gallardo et al., 2021, 2019; Villares et al., 2015; Gatchalian et al., 2013; Prieto et al., 2009; Trachana et al., 2007).

**Figure 1.**
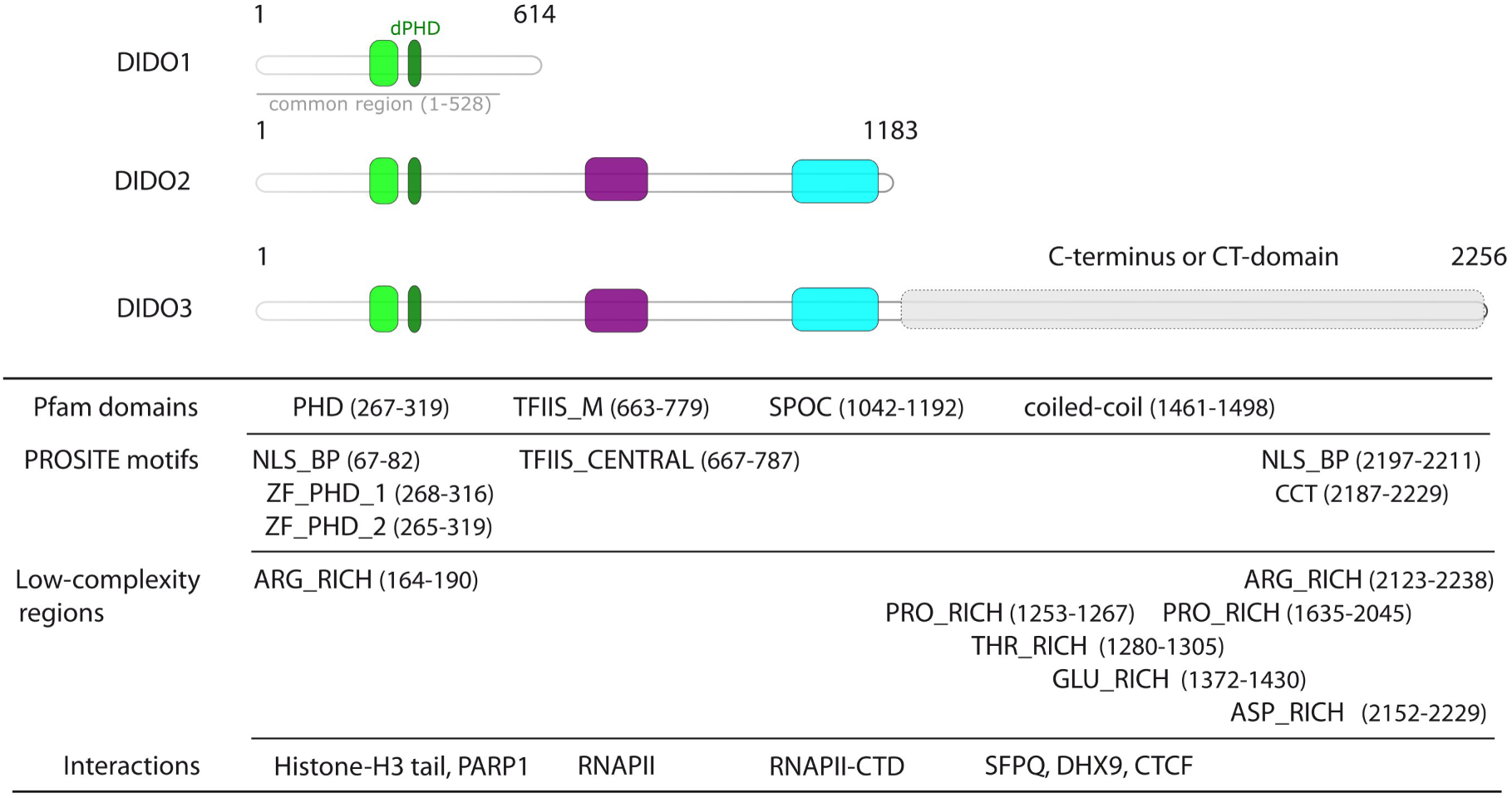
Scheme of the protein isoforms produced by alternative splicing of the *Dido1* gene. The amino acid start and end positions for the Pfam domains, PROSITE motifs and the low-complexity regions (indicated in parentheses) follow the numbering scheme of the mouse sequence (UniProtKB accession number: Q8C9B9). Experimentally determined protein interactions are illustrated at the bottom of the figure. The following abbreviations are used: PHD, PHD finger; dPHD, a predicted domain of approximately 25 amino acids located 67 residues downstream of the PHD domain (Rojas et al., 2005); TFIIS_M, transcription factor S-II central domain; SPOC, Spen paralogue and orthologue C-terminal domain; NLS_BP, bipartite nuclear localization signal; ZF_PHD_1 and ZF_PHD_2, zinc finger PHD-type; TFIIS_CENTRAL, TFIIS central domain; CCT, CCT domain; ARG_RICH, arginine-rich region; PRO_RICH, proline-rich region; THR_RICH, threonine-rich region; GLU_RICH, glutamic acid-rich region; ASP_RICH, aspartic acid-rich region. Annotations of the protein domains, motifs and low-complexity regions are available at http://pfam.xfam.org/ and https://prosite.expasy.org.

Histone marks like H3K4me3, H3K36me3 and H3K27ac are associated with genome activation (Martire & Banaszynski, 2020), with H3K4 and H3K36 methylation, and post-translational modifications (PTMs) at H3K27, contributing to facultative heterochromatin formation (Fang et al., 2021; Kasinath et al., 2021). The DIDO3 PHD domain acts as an epigenetic reader, binding preferentially to H3K4me3 (Tencer et al., 2017; Gatchalian et al., 2013). However, it remains unclear whether the N-terminal region of DIDO3 (DIDO3_1-528_, NT) binds to histone H3 peptides (which can be modified through acetylation and methylation at lysines 27 and 36), an issue that is important to understand how DIDO3 recognizes and interacts with distinct chromatin states.

Genetic alterations to *Dido1* have been linked to a range of different cancers (Zhang et al., 2021; Forghanifard et al., 2020; Li et al., 2020; Xiao et al., 2020; Assarnia et al., 2019; Thutkawkorapin et al., 2019; Yang et al., 2019; Lyu et al., 2017; Siemers et al., 2017; Berzoti-Coelho et al., 2016; Braig & Bosserhoff, 2013). However, the molecular mechanisms underlying these associations and the consequences of *Dido1* disruption are still poorly understood.

Consequently, here we present experimental and computational evidence that DIDO3 is associated with both RNAPII transcription and the regulation of chromatin structure, probably through its participation in multi-protein complexes. Our results suggest that DIDO3 may be associated with a feedback loop involving elements of RNAPII-associated complexes that contain DIDO3-binding sites. Furthermore, we highlight the association of DIDO3 with genomic regions linked to embryonic stem cell (ESC) self-renewal and differentiation, and genes implicated in cancer development.

## RESULTS

Our integrated computational analysis of DIDO3 binding sites (ChIP-seq, GSE85029), the transcriptomic data from the ΔE16 variant of DIDO3 in which exon 16 has been deleted (RNA-seq, GSE152346) and of DIDO3 protein-protein interactions (PPIs) indicate that the C-terminal (CT) domain of the protein is critical to drive the transcription of different subunits of heterooligomeric protein complexes (Figure 1). We identified a set of primary targets (TG1) for which DIDO3 exon 16 deletion affected both their expression and binding to DIDO3, as well as secondary targets (TG2) that only suffered altered expression (see Table 1; further details in Supplementary Table S1 and Supplementary Files 1 and 2). Interactions between DIDO3 and proteins encoded by genes containing DIDO3 binding sites were confirmed by proximity ligation assays, co-purification, co-immunoprecipitation and proteomics analyses (Table 2 and Supplementary Table S1). These proteins included the SFPQ splicing factor (Mora Gallardo et al., 2019), the heterogeneous nuclear ribonucleoprotein hnRNPU required for chromatin organization (Mora Gallardo et al., 2019), the helicase DHX9 that unwinds DNA and DNA:RNA hybrids (Fütterer et al., 2021), and the chromatin decondensation factor NCL (Mora Gallardo et al., 2019). Consequently, genes that encode proteins with DIDO3 binding sites and that interact with DIDO3 in heterooligomeric protein complexes defined a third group of targets (TG3). The identification of DIDO3 targets in TG1 and TG3 could reflect a possible feedback loop between protein subunits in RNAPII-associated complexes and genes encoding a DIDO3-binding site.

**Table 1.** Primary and secondary target genes that are expressed distinctly in DIDO3ΔE16 and wild-type ESCs. ^(a)^ Gene positions are indicated by the chromosome (Chr) number and strand direction (forward [+] or reverse [-]). ^(b)^ Each unique MACS2 identifier corresponds to a ChIP-seq peak from the HA-DIDO3 dataset (GSE85029). ^(c)^ Gene expression changes between DIDO3ΔE16 and wild-type embryonic stem cells, including log2 fold change (log_2_FC) and adjusted p-values (false discovery rate, FDR) that were calculated from the RNA-seq data using edgeR. ^(d)^ UniProtKB annotations are provided (www.uniprot.org): a hyphen (-) indicates that no ChIP-seq peak was detected.

| *Gene* | *Chr* <sup>(a)</sup> | *ChIP-seq* <sup>(b)</sup> | *RNA-seq* <sup>(c)</sup> | *Description* <sup>(d)</sup> |
| --- | --- | --- | --- | --- |
| **TG1 group** |  |  |  |  |
| *Carm1* | Chr9 (+) | #2735 | log<sub>2</sub>FC = -0.81<br>FDR = 1.73e-03 | Histone methyltransferase (H3R17me2) |
| *Setd7* | Chr3 (-) | #1718 | log<sub>2</sub>FC = -0.86<br>FDR = 1.16e-02 | Histone methyltransferase (H3K4me1) |
| *Dido1* | Chr2 (-) | #1664 to #1671 | log<sub>2</sub>FC = -4.81<br>FDR = 1.21e-11 | H3-tail reader, RNA pol II-associated |
| *Hmga1* | Chr17 (+) | #1174, #1175 | log<sub>2</sub>FC = -1.56<br>FDR = 2.77e-10 | Nucleosome phasing |
| *Dhx16* | Chr17 (+) | #1220, #1221 | log<sub>2</sub>FC = -1.07<br>FDR = 6.49e-04 | mRNA splicing |
| *Sall1* | Chr8 (-) | #2682 | log<sub>2</sub>FC = -1.13<br>FDR = 6.27e-04 | Transcription repressor (pluripotency and differentiation) |
| *Mau2* | Chr8 (-) | #2626, #2627 | log<sub>2</sub>FC = -3.03<br>FDR = 3.24e-16 | Cohesin-loading complex (chromatid cohesion factor) |
| *Aebp2* | Chr6 (+) | #2299, #2300 | log<sub>2</sub>FC = 2.12<br>FDR = 8.72e-08 | Polycomb accessory protein (PRC2.2) |
| *Epop* | Chr11 (-) | #509, #508 | log<sub>2</sub>FC = 1.00<br>FDR = 1.44e-08 | Polycomb accessory protein (PRC2.1) |
| *Trim28* | Chr7 (+) | #2336 | log<sub>2</sub>FC = -0.81<br>FDR = 1.73e-03 | Scaffold protein for the heterochromatin protein 1 (CBX5/HP1A) |
| *Set1b/Setd1b* | Chr5 (+) | #2148 | log<sub>2</sub>FC = 1.01<br>FDR = 2.7e-03 | Set1C/COMPASS complex protein, methyltransferase activity (H3K4me2/3) |
| *Gltscr1l/Bicral* | Chr17 (-) | #1242 | log<sub>2</sub>FC = 0.88<br>FDR = 3.35e-04 | SWI/SNF complex (GBAF) |
| *Smarcad1* | Chr6 (+) | #2225 | log<sub>2</sub>FC = 0.92<br>FDR = 2.02e-02 | SWI/SNF-like remodeler |
| *Smarcc1* | Chr9 (+) | #2838 | log<sub>2</sub>FC = -1.25<br>FDR = 1.51e-04 | SWI/SNF complex (GBAF, nBAF, npBAF) |
| **TG2 group** |  |  |  |  |
| *Bhlhe40* | Chr6 (+) | - | log<sub>2</sub>FC = -1.44<br>FDR = 3.46e-07 | Novel CTCF binding partner |
| *Birc5/Survivin* | Chr11 (+) | - | log<sub>2</sub>FC = 7.76<br>FDR = 3.35e-03 | Recruitment of chromosome passage protein complex (CPC) to centromeres |
| *Cenpb* | Chr2 (-) | - | log<sub>2</sub>FC = 0.81<br>FDR = 2.19e-03 | Centromere-associated protein |
| *Cenpm* | Chr15 (-) | - | log<sub>2</sub>FC = 0.73<br>FDR = 1.21e-02 | Centromere-associated protein |
| *Foxd3* | Chr4 (+) | - | log<sub>2</sub>FC = 1.5<br>FDR = 1.85e-05 | Maintenance of pluripotent cells |
| *Mapk11* | Chr15 (-) | - | log<sub>2</sub>FC = -0.95<br>FDR = 1.82e-02 | Phosphorylate histone H3 and the nucleosomal protein HMGN1 |
| *Pcl1/Phf1* | Chr17 (+) | - | log<sub>2</sub>FC = -1.71<br>FDR = 4.2e-02 | Polycomb accessory protein (PRC2.1), H3K36me3 reader |
| *Setd5* | Chr6 (+) | - | log<sub>2</sub>FC = -4.17<br>FDR = 3.28e-26 | Histone methyltransferase (H3K36me3) |
| *Sugp1* | Chr8 (+) | - | log<sub>2</sub>FC = -1.07<br>FDR = 2.46e-02 | mRNA splicing |
| *Zfp462* | Chr4 (+) | - | log<sub>2</sub>FC = -0.82<br>FDR = 1.94e-02 | Nuclear factor (chromatin structure, pluripotency and differentiation), regulates SOX2, POU5F1/OCT4, and NANOG |

**Table 2.**
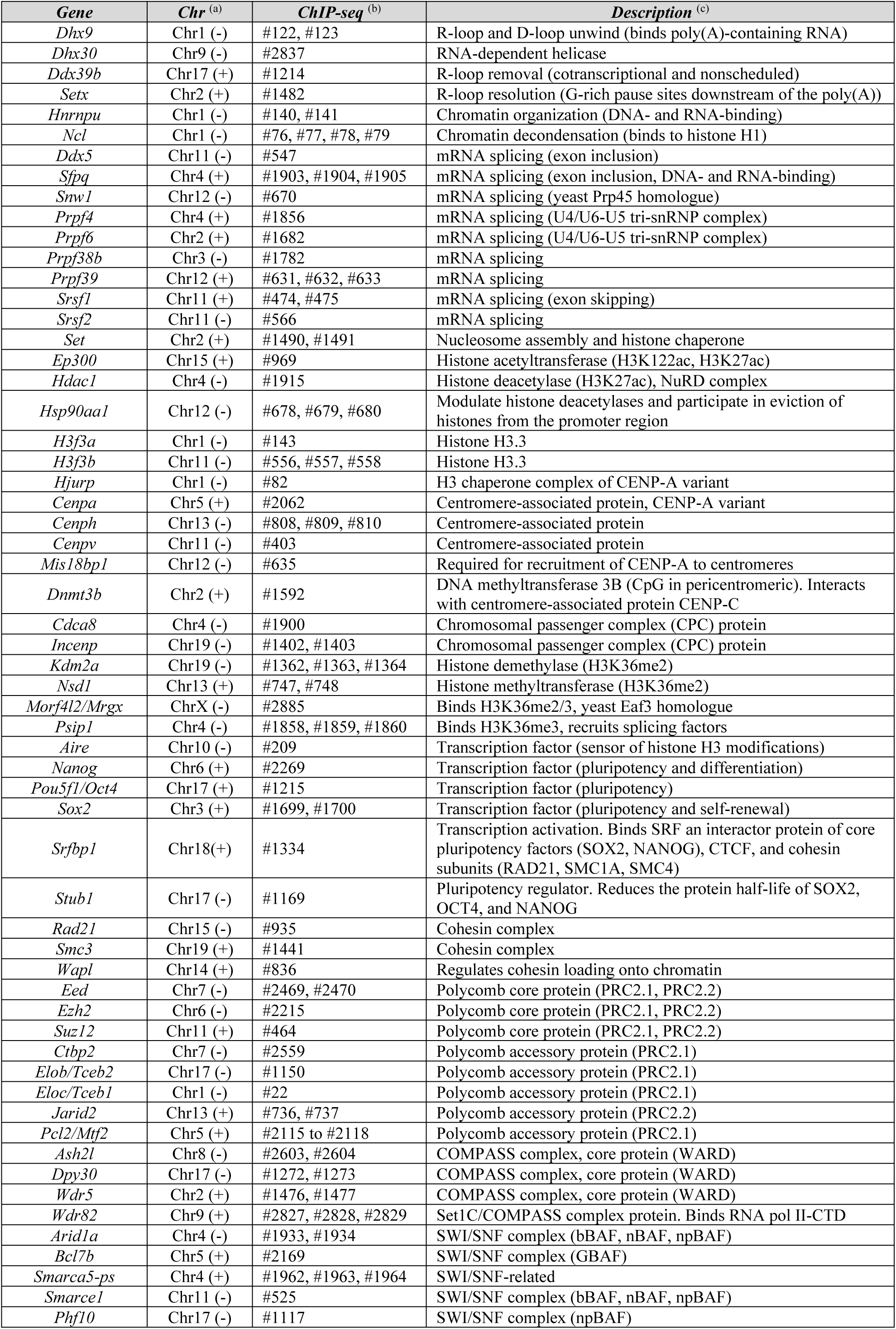
Genes with DIDO3 binding sites that could interact with DIDO3 and their proteins. ^(a)^ Gene position is represented by the chromosome (Chr) number and strand directions in parentheses as forward (+) or reverse (-). ^(b)^ Unique MACS2 identifiers correspond to the ChIP-seq peak from the HA-DIDO3 dataset (GSE85029). ^(c)^ UniProtKB annotations are provided (www.uniprot.org).

### DIDO3 binding and co-localization

ChIP-seq experiments with wild-type (WT) DIDO3 ESCs (GSE85029) indicated that DIDO3 and RNAPII co-localize extensively in the mouse genome, with 79% of DIDO3 binding sites (2269 of 2888) overlapping with that for the largest subunit RNAPII, RPB1 (Figure 2A and Supplementary Table S2). This finding is consistent with studies of a RNAPII ChIP-seq dataset (GSM2645517) that confirmed DIDO3 and RNAPII binding at the *Dido1* locus, particularly at the proximal promoter and an internal binding site (Fütterer et al., 2017). Genome-wide computational analysis of DIDO3, epigenetic marks and chromatin regulators further revealed a significant and unexpected overlap of DIDO3 binding sites with H3K36me3 (69%, 2002 intersect sites), CTCF (13%, 360 intersect sites) and PRC2 (8%, 242 intersect sites: Figure 2A and Supplementary Table S2). Aggregate peak analysis of ChIP-seq and Hi-C data (Galan et al., 2022; Rao et al., 2014) indicated that DIDO3 binding sites are situated close to RNAPII, H3K36me3, H3K4me3 and H3K27ac sites in the 3D chromatin structure of ESCs (Figure 2B), with these interactions being more prominent in genes downregulated in DIDO3ΔE16 datasets than in those upregulated genes (Figure 2B, last two rows).

**Figure 2.**
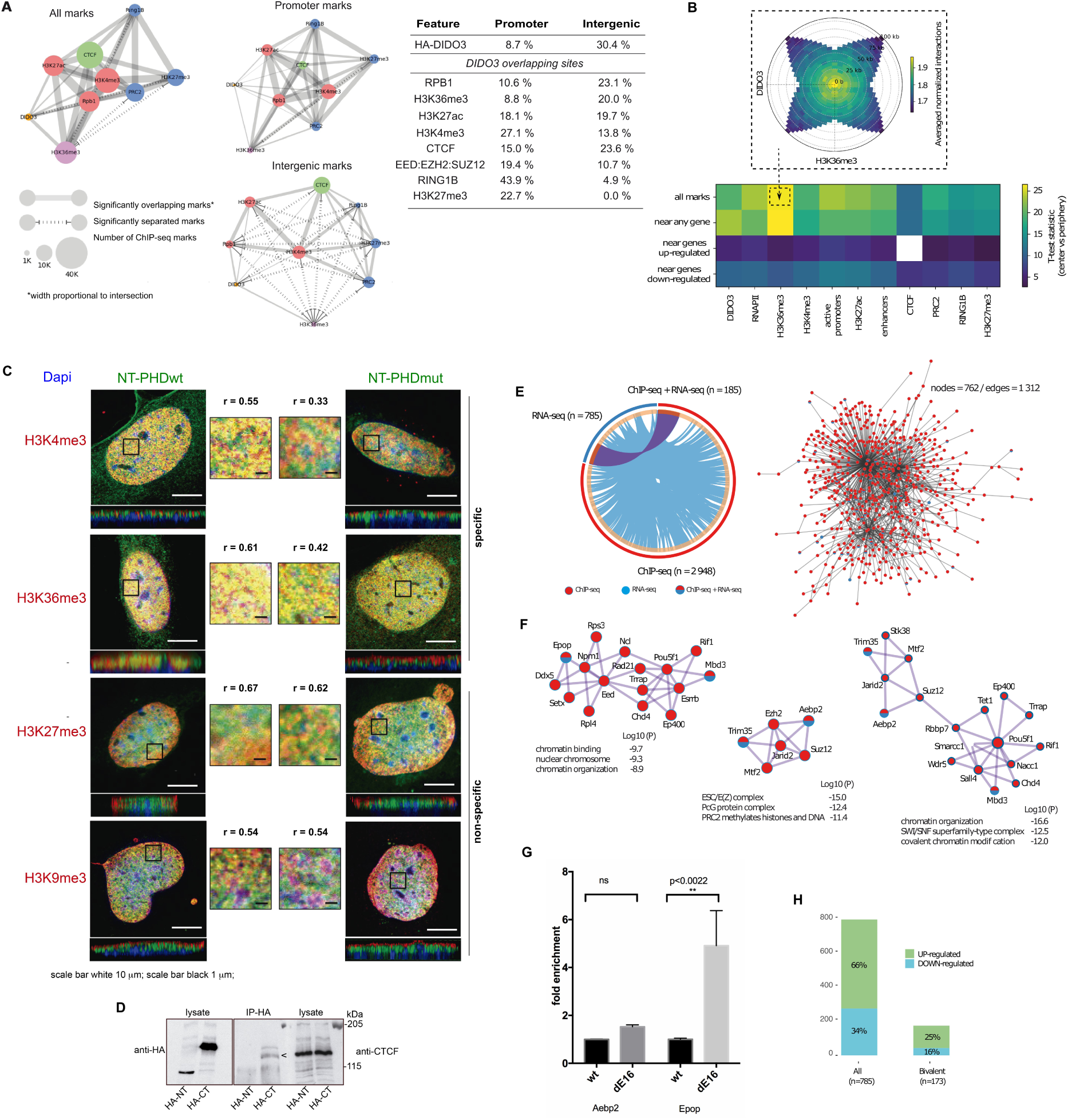
DIDO3 binding site intersection with RNAPII, CTCF, Polycomb and histone H3 modifications. (A) Distribution of overlapping peaks among DIDO3, RNAPII, CTCF, Polycomb and H3 modifications across chromosomes (left), and the percentage of these overlapping peaks located in promoters and intergenic regions (right). (B) Example of an Aggregate Peak Analysis (APA) in polar coordinates showing the average interaction enrichment between DIDO3 and H3K36me3 coordinates (inset). Co-localization of DIDO3 binding sites with other ChIP-seq marks in the 3D chromatin structure of mouse embryonic stem cells (ESCs) is shown in four rows: the first row displays DIDO3 marks against other marks; the second row shows interactions between marks near any gene; the third row presents interactions near genes upregulated in DIDO3ΔE16; and the fourth row shows interactions near genes downregulated in DIDO3ΔE16. Proximity between the gene and mark is defined as being within the same 5 kb bin. (C) Localization of wild-type (NT-PHDwt) or mutant (NT-PHDmut) N-terminal PHD domains and histone H3-specific trimethylation on several distinct lysine residues. The upper row shows co-localization of NT and H3K4me3 (left), which is dampened by the PHD mutation (right). The second row demonstrates strong co-localization of NT with H3K36me3 (left), with a slight decrease observed upon PHD mutation (right). The third row shows non-specific co-localization with H3K27me3 (right) and the fourth row shows non-specific co-localization with H3K9me3 (left). Representative optical sections and a portion of their 3D model are shown. Scale bars represent 10 μm for whole cells and 1 μm for the amplified areas. The Pearson’s coefficient for co-localization was calculated as described in Appendix 1. (D) Interaction between CTCF and the DIDO3 CT region. Lysates from ΔCT stably transfected ESCs expressing either HA-tagged N-terminal (HA-NT) or C-terminal (HA-CT) DIDO3 fragments were immunoprecipitated with an anti-HA rabbit polyclonal antiserum (ab9110, Abcam). Co-immunoprecipitation of CTCF was detected in samples overexpressing HA-CT by immunoblotting with an anti-CTCF polyclonal antiserum (07-729; Millipore). (E) The left panel displays a Circos plot illustrating the overlap between genes identified by ChIP-seq and RNA-seq analyses, while the right panel shows the protein-protein interaction (PPI) network of these overlapping genes. (F) Densely connected sub-networks identified with the MCODE algorithm highlighting clusters of interacting proteins within the PPI network. (G) Quantitative RT-PCR analysis of Aebp2 and Epop expression. (H) Bivalent genes upregulated and downregulated in DIDO3ΔE16 ESCs.

### DIDO3 co-localizes with histone H3K4me3 and H3K36me3

To test whether DIDO proteins interact with different histone H3 lysine trimethylation marks, we examined the co-localization of DIDO3 with H3K4me3, H3K9me3, H3K27me3 and H3K36me3 in mouse embryonic fibroblasts (MEFs) by confocal microscopy (Figure 2C). HA-tagged DIDO3 NT constructs containing either a WT PHD domain or a PHD domain carrying the C285A/H290A mutation that disrupts H3K4me3 binding were overexpressed in E6/E7 immortalized MEFs from ΔNT mutant cells (Gatchalian et al., 2013). As expected, the fluorescent signals of NT and H3K4me3 co-localized with the WT PHD domain, but this signal was dampened when this PHD domain was mutated (Fig 2C upper panel). Similarly, strong co-localization of NT with H3K36me3 was observed, which was only mildly dampened by the PHD mutation (Figure 2C, second panel), suggesting that regions of the DIDO3 NT domain other than the PHD domain also contribute to the H3K36me3 interaction. Conversely, co-localization between NT and either H3K27me3 or H3K9me3 was unaffected by the PHD mutation, indicating that these modifications do not specifically interact with the PHD domain (Figure 2C; see also Supplementary Figure S1A).

### The association of DIDO3 with chromatin insulation and long-distance contacts

Higher-order chromatin organization is controlled by several proteins like RNAPII, CTCF, cohesin, and PRC2. RNAPII is fundamental for both chromatin compartmentalization and chromatin loop establishment following mitosis (Zhang et al., 2021), while CTCF and cohesin play critical roles in chromatin loop formation and the genome-wide insulation of topologically associated domains (TAD) (Jerković et al., 2020). CTCF also interacts with and recruits the RPB1 subunit of RNAPII to its target sites in the genome (Chernukhin et al., 2007). Our ChIP-seq and RNA-seq analysis identified genes encoding the cohesin subunits RAD21 and SMC3, as well as the cohesin DNA release factor WAPL in TG3 (Table 2), and the cohesin-loading factor MAU2 in TG1 (Table 1). Three of these proteins (MAU2, RAD21 and SMC3) interact through the NIPBL-MAU2 heterodimer and the cohesin complex (composed of SMC1A, SMC3, RAD21 and STAG1). Also, BHLHE40 was identified as a novel CTCF (TG2), and the loss of BHLHE40 in HeLa-S3 cells reduces CTCF binding to its recognition sites and disrupts CTCF-mediated long-range chromatin interactions (Hu et al., 2020). Interestingly, the *Mau2* and *Bhlhe40* genes were downregulated in RNA-seq data from DIDO3ΔE16 mutant cells (Table 1). The interaction of CTCF with DIDO3 was assessed in DIDO3ΔE16 ESCs transfected with HA-tagged NT or CT domains and in western blots of the proteins immunoprecipitated with anti-HA, CTCF was seen to interact with the CT-domain but not the NT domain of DIDO3 (Figure 2D).

Our meta-analysis of gene expression in DIDO3ΔE16 ESCs (GSE152346), DIDO3 binding sites (GSE85029) and PPIs suggests that DIDO3 is associated with genomic regions involved in long-distance chromatin contacts mediated by Polycomb complexes (Tables 1 and 2, and Figure 2E-G). Gene repression by the Polycomb repressive complex 2 (PRC2) is mediated by two functional assemblies, PRC2.1 and PRC2.2, which share common core subunits but utilize distinct accessory subunits. PRC2.1 preferentially maintains gene repression, while PRC2.2 mainly mediates the de novo repression of active genes (Petracovici & Bonasio, 2021). The core PRC2 subunits EED, EZH2 and SUZ12 (TG3) are found in densely connected regions of the PPI network (Table 2 and Figure 2F). Several PRC2 accessory proteins were found in TG1 or TG3, including AEBP2, CTBP2, ELOB (TCEB2), ELOC (TCEB1), EPOP, JARID2, PHF1 and PCL2 (MTF2: Tables 1 and 2), and these were located in densely connected regions of the PPI network (Figure 2F). AEBP2 is a specific accessory protein of PRC2.2 that acts as a DNA-binding transcriptional repressor for RNAPII. By contrast, EPOP serves as a scaffold between the PRC2/EED-EZH2 complex and the elongin BC (ELOB-ELOC) complex, playing a key role in bivalent chromatin regions that possess both active and repressor properties (Mas et al., 2018). Recent evidence suggests that EPOP disrupts the PRC2.1 dimer and weakens its association with chromatin (Gong et al., 2024). Notably, *Aebp2* and *Epop* were overexpressed in DIDO3ΔE16 mutant cells (Table 1), a finding confirmed by qRT-PCR (Figure 2G).

Because PRC2 activity alters H3K27 methylation (Petracovici & Bonasio, 2021), the global H3K27 methylation was assessed at the onset of differentiation of WT and DIDO3ΔE16 ESCs. Histone lysates from ESCs and early embryonic bodies (EBs) of both genotypes were probed for mono-, di- and tri-methylated, as well as total H3K27, but no differences between WT and mutant cells were observed (Supplementary Figure S2). Hence, any differences in H3K27 methylation may be restricted to specific target genes. The *Phf1* and *Mtf2* genes encode two H3K36me3 readers that recruit PRC2 to H3K36me3-containing chromatin regions (Cai et al., 2013). *Phf1* (TG2) appears to be regulated by DIDO3, as it is downregulated in DIDO3ΔE16 ESCs, while *Mtf2* expression was unaltered but it did contain a DIDO3 binding site (TG3: Tables 1 and 2).

Genes regulating ESC development and differentiation are enriched with bivalent promoters, known targets of Polycomb, and they undergo transcriptional switching upon differentiation (Kumar et al., 2021). Bivalency maintains weak gene expression while keeping them poised for rapid activation. The RNA-seq analysis of WT and DIDO3ΔE16 mutant ESCs indicated that approximately 25% of the upregulated genes and 16% of the downregulated genes in the latter (Figure 2H) were *bona fide* bivalent genes, as described in mouse ESCs (Mas et al., 2018). Moreover, bivalent genes were significantly more likely to be upregulated relative to non-bivalent genes in DIDO3ΔE16 ESCs (Fisher’s test p=0.005; Figure 2H). Among the *bona fide* bivalent genes, *Bhlhe40*, *Foxj2*, *Mapk11*, *Otx2* and *Setd7* were downregulated, while *Pus3* and *Foxd3* were overexpressed (Table 1 and Supplementary Table S1). The DIDO3 binding to PRC2 core subunits in WT ESCs, along with transcriptional dysregulation of PRC2 accessory proteins and bivalent genes in DIDO3ΔE16 mutant cells, suggests that DIDO3 may influence 3D chromatin organization.

When the relationship between DIDO3 binding near transcription start sites (TSS) of up- and downregulated genes was assessed in DIDO3ΔE16 ESCs (Supplementary Figure S3, left), the frequency of DIDO3 binding sites around the TSS (from -50kb upstream to +50kb downstream) was similarly distributed in both. DIDO3 binding to TSS was above random at up- or down-regulated genes in DIDO3ΔE16, although this enhanced specificity was comparable to that for other proteins where ChIP-seq data is available (Supplementary Figure S3, right). Thus, DIDO3 does not appear to be the main regulator of gene expression at these sites but rather, it may be associated with RNAPII-mediated transcriptional activation and repression, consistent with a recent review of domain acquisition in the TFIIS transcription factor (TF) family (Matveyeva et al., 2025).

CTCF was recently assigned dual functions, as an insulator for somatic gene silencing and as a chromatin remodeller for pluripotency gene activation during reprogramming (Song et al., 2022). DIDO3 interacts with CTCF (Figure 2D), and it has essential functions in ESC differentiation and somatic cell reprogramming (Fütterer et al., 2021). The chromatin regulators *Setd7* (TG1), *Bhlhe40* (TG2) and *Srfbp1* (TG3) exemplify the proposed feedback loop (Tables 1 and 2), with SETD7 interacting with the EED and SUZ12 PRC2 subunits, among other proteins, while BHLHE40 interacts with CTCF. SRFBP1 binds to SRF that interacts with core architectural proteins mediating chromatin loops and topologically associating domains (TADs). These include CTCF, and the cohesin subunits RAD21, SMC1A and SMC4 (Tsaytler et al., 2024). The bivalent genes *Setd7* and *Bhlhe40* were downregulated in DIDO3ΔE16 mutant cells (Table 1).

Taken together, these findings are consistent with the possibility that DIDO3 is associated with regions involved in 3D chromatin organization, which should be confirmed in Hi-C experiments on DIDO3ΔE16 mutant cells. Nevertheless, various genes in TG1 (*Dido1*, *Hmga1*, *Klf2*, *Smarcc1*, *Smarcad1*, *Pus3*) have been associated with intra-chromosomal CTCF-CTCF interactions in ESCs using an independent CTCF ChIA-PET dataset (Handoko et al., 2011).

### DIDO3 marks are enriched at TAD boundaries and in active A-compartments

In the light of the interactions among DIDO3, RNAPII, CTCF and PRC2, the relationships between DIDO3 binding sites, chromatin loops (CTCF and Polycomb) and TAD boundaries were studied. The combined analysis of ChIP-seq (GSE85029, GSM2645517, GSM2645511, GSM2645495, GSM2645513, GSM2645515, GSM2645497, GSE120393, GSM2645509) and Hi-C data in ESCs (GSE99530) showed enrichment of DIDO3, RNAPII, H3K36me3, H3K4me3, H3K27ac and CTCF binding sites inside the chromatin loops (Figure 3A). DIDO3 marks showed the clearest enrichment inside loops, between loop-anchor points (positive coordinates in Figure 3A). While DIDO3 previously overlapped other marks, here it could be differentiated from CTCF sites and from RNAPII to a lesser extent. Six classes of chromatin loops were defined (all, Polycomb, enhancer-promoter, CTCF, CTCF+enhancer-promoter and CTCF alone) and enrichment of DIDO3, RNAPII, H3K36me3, H3K4me3 and H3K27ac was assessed in each (Figure 3B). Consequently, DIDO3 and RNAPII binding sites were enriched in “CTCF only” but not in “Polycomb” loops (Figure 3B), suggesting that DIDO3 is physically pushed toward the inside of chromatin loops by CTCF occupancy or that of its associated proteins. In the context of TAD boundaries (Figure 3C), the expected enrichment in CTCF was observed, but also that of RNAPII and histone marks related to transcriptional activity (H3K4me3, H3K27ac and H3K36me3). Of the proteins and histone marks measured, DIDO3 is among those most enriched at TAD boundaries. The distribution of DIDO3 marks in chromatin compartments was also analyzed and as suggested previously, DIDO3 marks were enriched in transcriptionally active chromatin (A compartments, Figure 3D). Genes that were activated or repressed in DIDO3ΔE16 ESCs were similarly distributed when analyzing genomic bins (100 kb resolution, Figure 3E). Downregulated genes were slightly more frequent than upregulated genes in A compartments, although this modest difference cannot be directly attributed to the presence of DIDO3 marks.

**Figure 3.**
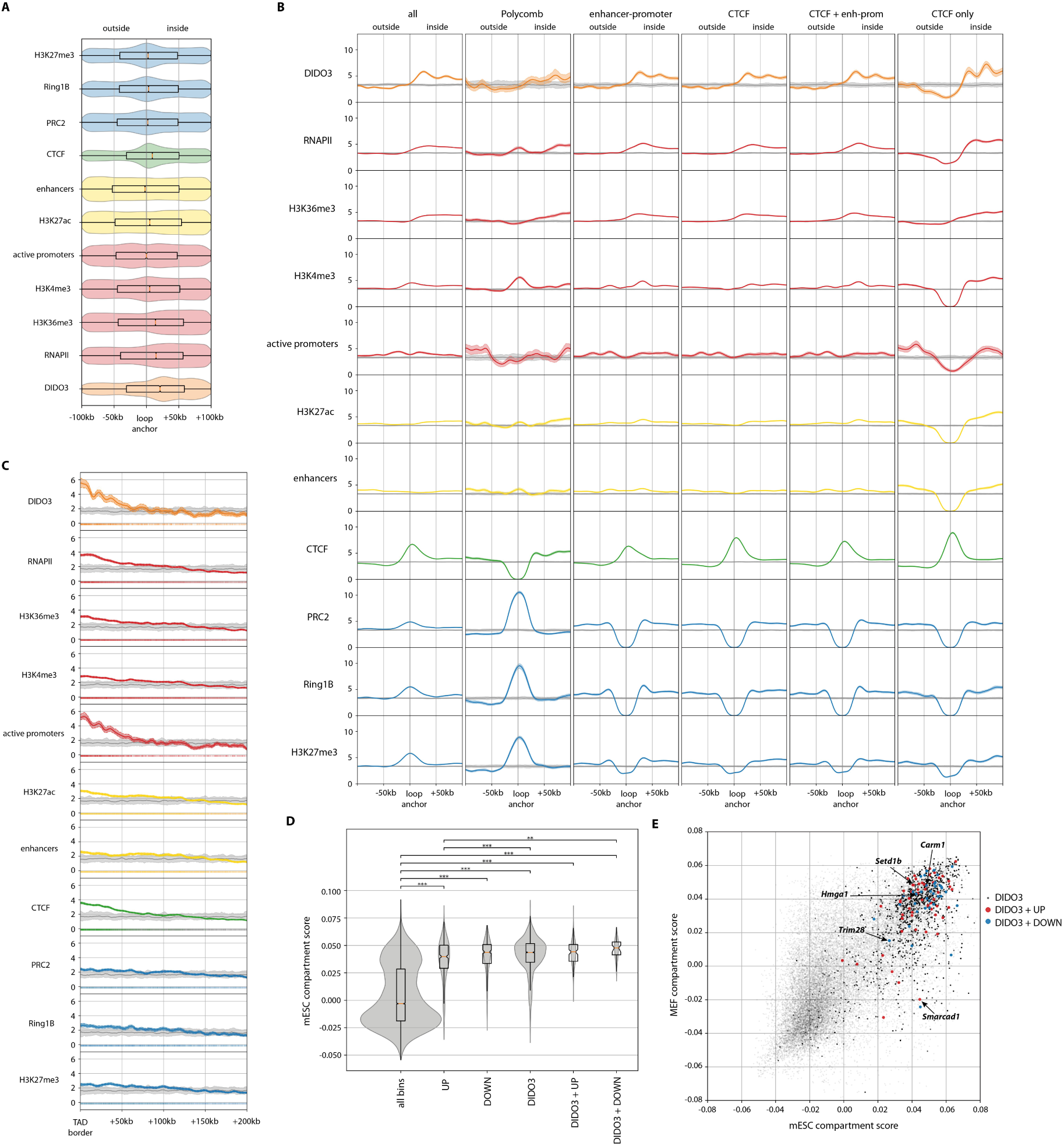
DIDO3 is associated with chromatin structure. *Active promoter* category, promoter regions with both H3K27ac and H3K4me3 marks; *enhancers,* regions enriched in H3K27ac at least 40 kb from any gene position of ChIP-seq marks relative to loop anchor points. Positive coordinates represent the inside of loops (between loop anchor points). (A) Overall position of ChIP-seq marks around loop anchor points. Notches in the boxplots represent 95% confidence of the median after 1,000 bootstraps. (B) Enrichment of ChIP-seq marks around loop anchor points with a 100 bootstrap error band. In grey, the result of 100 randomizations of the ChIP-seq mark coordinates: *all* represents any loop detected by the software Mustache; *Enhancer-promoter*, loops with anchor points away from PRC2 or Ring1B (>20 kb) and close to H3K27ac or H3K4me3 (<40 kb); *CTCF*, loops with anchor points away from PRC2 or Ring1B (>20 kb) and close to CTCF marks (<20 kb); *CTCF + enh-prom*, loops with anchor points away from PRC2 or Ring1B (>20 kb), close to H3K27ac or H3K4me3 (<40 kb) and close to CTCF marks (<20 kb); *CTCF only*, loops with anchor points away from PRC2 or Ring1B or H3K27ac or H3K4me3 (> 20 kb), and close to CTCF marks (< 20 kb). (C) As in (A), with TAD borders detected by TADbit. (D) Distribution of compartment scores in all 100 kb genomic bins (left), bins containing: upregulated genes in DIDO3ΔE16; downregulated genes in DIDO3ΔE16; bins with DIDO3 ChIP-seq marks; bins with the mark and upregulated genes; and finally (right), bins with the mark and downregulated genes. (E) Changes in the A/B compartment score in mESCs and MEFs: the bins in black indicate genes with a DIDO3 mark in mESCs; in red, genes with DIDO3 marks and a significant increase in expression in mESCs; and in blue, genes with DIDO3 mark and a significant decrease in expression in mESCs.

### DIDO3 is a candidate “reader” of H3K36me3 and may be associated with regions involved in pre-mRNA splicing and facultative heterochromatin

PHD domains read histone states in a sequence-dependent manner that is positively or negatively modulated by lysine and arginine methylation, and by lysine acetylation (Sanchez & Zhou, 2011). DIDO3 binds preferentially to two distinct regions (Figure 2A), on the one hand it binds to the gene body and intergenic regions, overlapping with RNAPII and H3K36me3, while on the other hand DIDO3 binds to enhancers and promoters, overlapping with H3K4me3, H3K27ac and CTCF. When the fluorescent signals of DIDO3 NT, H3K4me3 and H3K36me3 were examined (Figure 2C) H3K36me3 was separate from any other mark we studied in the intergenic regions (Figure 2A and Supplementary Table S2), indicating it is not a secondary target of DIDO3. The marked overlap between DIDO3 and H3K36me3 binding sites (69%, 2002 intersect sites) in the mouse genome and its isolation from any other mark in intergenic regions, indicate that this interaction is highly specific, prompting closer study of the amino acid sequence of the DIDO3 NT (PHD and dPHD domains).

The DIDO3 PHD domain contains most of the key residues in the H3K36me3-binding pocket of ECM5 PHD (Shi et al., 2007). Based on the alignment and residue location in the DIDO3 PHD crystallographic structure, H3K36me3 binding could involve the Y267 (region III), H274 (region IV), M279 and C281 (region II), W288 (region I) at the equivalent positions to ECM5-Y1240, ECM5-E1247, ECM5-M1252, ECM5-E1254 and ECM5-W1261, respectively. Other combinations of residues at these positions in the H3K36me3-binding pocket may also contribute to binding (Figure 4A, top panel). The AlphaFold3 structural prediction of the DIDO3 PHD–H3K36me3 interaction reveals that H3K4 and H3K36 occupy similar positions within the PHD binding pocket (Figure 4B). The predicted dPHD domain (Rojas et al., 2005) has a Cys4His motif instead of the canonical Cys4HisCys3 in PHD (Figure 4A, bottom panel) and hence, the three-dimensional orientation of the PHD and dPHD domains was also examined using AlphaFold3. As AlphaFold predicts the two domains to be spatially separated, dPHD is unlikely to participate in H3-tail recognition by PHD (Figure 4C).

**Figure 4.**
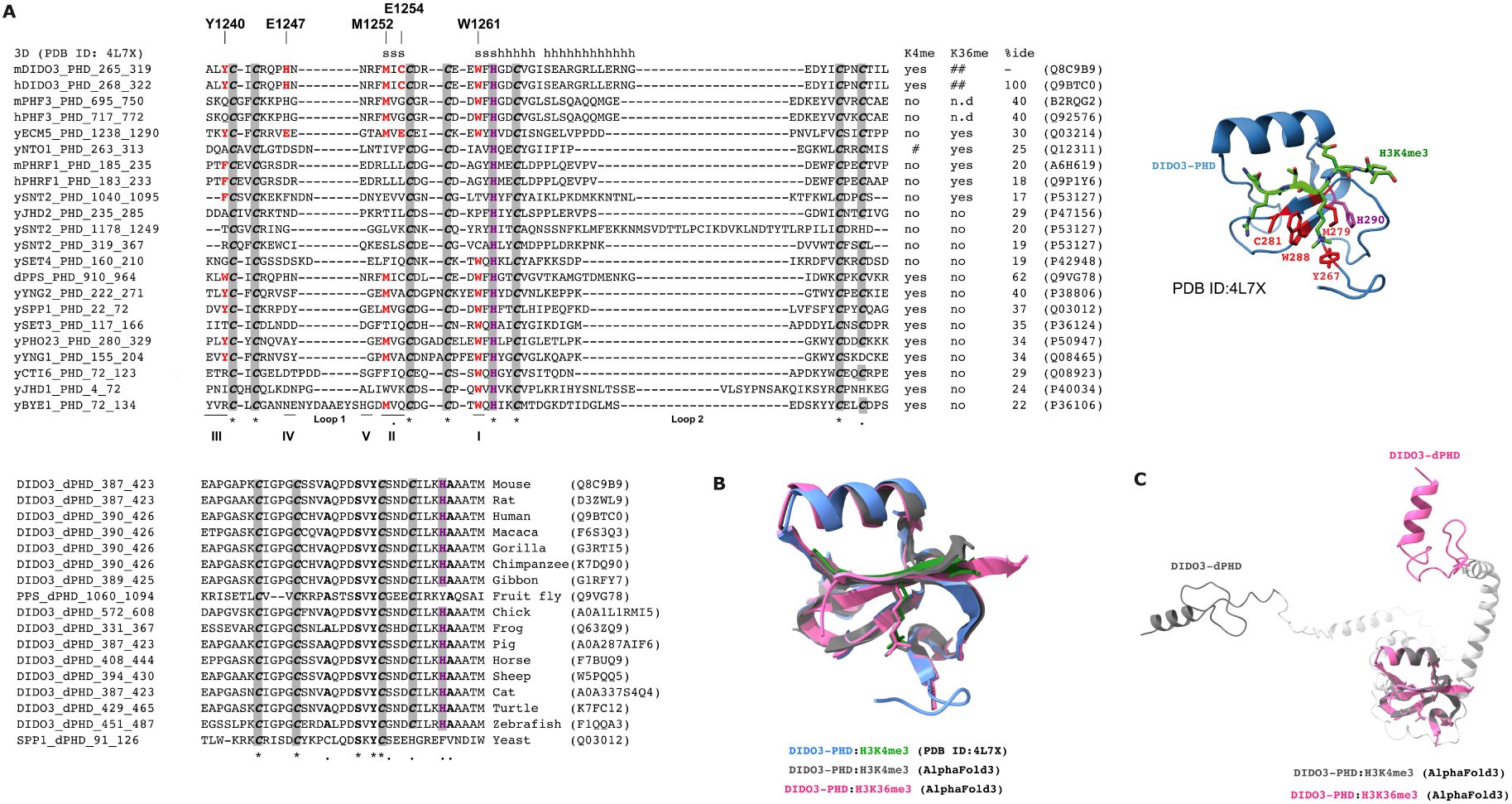
The DIDO3 PHD domain shares most key residues in the peptide-binding pocket for H3K4me3 and H3K36me3. (A) Sequence comparison of yeast and mouse PHD (top) and dPHD domains (bottom), with the regions involved in ligand recognition labeled I through V (Sanchez & Zhou, 2011). The DIDO3 PHD crystallographic structure and the positions of the key residues in the peptide-binding pocket are shown on the right. (B) Structural alignment of the crystallographic structure and AlphaFold3 predictions for PHD-H3K4me3 and PHD-H3K36me3 interactions. (C) Structural alignment of the AlphaFold3 prediction for PHD and dPHD domains.

Surface plasmon resonance imaging (SPRi) was used to quantify the binding of the DIDO3 PHD domain to 11-mer histone H3 tail peptides (Supplementary Figure S1B). The PHD domain bound H3K4me3 (KD = 25.5 nM), H3K9me3 (KD = 22.2 nM) and H3K27me3 (KD = 26.5 nM) with comparable affinities, whereas binding to H3K36me3 was modestly weaker (KD = 32.1 nM). Although these affinities differ from those reported for other PHD domains, all the interactions lie within the low nanomolar range (Supplementary Figure S1B and C). Notably, the dissociation rate constant (kd) that reflects the stability of the complex was nearly identical for H3K4me3 (23.8 x 10^-4^ s^-1^) and H3K36me3 (24.0 x 10^-4^ s^-1^), confirming that the DIDO3 PHD domain recognizes both H3K4me3 and H3K36me3 *in vivo* and *in vitro* (Figure 2C and Supplementary Figure S1B).

The co-existence of H3K4me3 and H3K36me3 at the same nucleosome has been shown through combinatorial-iChIP experiments in *S*. *cerevisiae* (Sadeh et al., 2016). The accumulation of both H3K4me3 and H3K36me3 in the same nucleosome is more likely to occur at highly expressed genes that experience repeated cycles of RNAPII passage (Sadeh et al., 2016). Thus, the presence of DIDO3 at sites marked by H3K36me3 in intergenic regions is intriguing, as H3K36me3 in these regions is associated with facultative heterochromatin or the DNA damage response (Cai et al., 2013; Li et al., 2013). Overlapping DIDO3 and H3K36me3 binding in gene body regions is consistent with the DIDO3ΔE16 mutant producing splicing defects (Mora Gallardo et al., 2019). H3K36me3 is required for efficient pre-mRNA splicing (Leung et al., 2019), and other genes necessary for pre-mRNA splicing and related to spliceosome assembly were identified as TG3 targets (Prpf4, Prpf6, Prpf38b, Prpf39, Srsf1, Srsf2: Table 2).

### DIDO3 and Transcription Factor interplay across promoters

Despite the absence of experimental evidence for direct DIDO3-DNA interactions, ChIP-seq data identified 2888 high-confidence DIDO3-bound genomic regions that were associated with the nearest TSS and flanking genes (up to 5 kb) on both strands (Supplementary File 2). We compared all 209 nucleotide sequences from DIDO3-bound promoter and proximal regions (3′ of the promoter) identified by ChIP-seq against known TF binding sites (mouse and human) annotated in HOCOMOCO v11, JASPAR 2022, SwissRegulon and a collection of ESCs TFs (Chen et al., 2008). MEME-ChIP analysis identified a 14 nucleotide motif in 39 out of 209 sequences, with an E-value of 8.4 x 10^-36^ (Table 3, see Supplementary Figure S4A for a comparison of this motif with known TF DNA-binding motifs). Some AT-rich DIDO3-associated promoter regions share sequence similarity with TF binding sites known to participate in differentiation-related regulatory loops, such as the BPTF nucleosome-remodeling factor subunit that interacts with H3K4me3, FOXJ3 (Forkhead box protein J3), ONECUT1 (hepatocyte nuclear factor 6) and ZNF384 (zinc finger protein 384) binding sites (Table 3).

**Table 3.** Overlap of specific sites for known TFs in DIDO3-bound promoter and proximal regions. This table lists vertebrate (e.g. *Mus musculus* and *Homo sapiens*) transcription factors (TFs) annotated in the target databases that have binding sites with partially overlapping nucleotide sequences in the DIDO3-bound promoter and in the proximal regions located 3′ of the promoter: ‘Database’ refers to the source database containing known TF motifs (e.g. HOCOMOCO v11, JASPAR 2022, SwissRegulon, and Chen2008, a collection of ESC TFs); ‘TF_ID’ refers to the identifier or name of the known transcription factors; ‘Class’ indicates the classification of the TF, such as its structural family or functional group; the ‘MEME-ChIP E-value’ represents the statistical significance of motifs (with E-value ≤ 0.05) identified by the programs MEME, STREME and CentriMo, all executed with default parameters unless otherwise specified; ‘Overlap’ is the number of motif columns that overlap in the optimal alignment; ‘Offset’ is the number of positions by which the query motif is shifted relative to the target motif in the optimal alignment, a positive value indicates the query is shifted downstream relative to the target motif; ‘p-value’ is the probability that a random motif of the same width as the target motif would achieve an optimal alignment with a match score equal to or greater than that observed for the target motif; ‘Tomtom E-value’ indicates the statistical significance of motif matches, with values of 1 or smaller considered significant; The ‘q-value’ represents the minimum false discovery rate (FDR) at which a match is deemed significant. Tomtom estimates p-values by sampling motif columns from the set of target motifs and calculates q-values from all match p-values using the Benjamini & Hochberg method (Benjamini & Hochberg, 1995). Additional motif logo information is provided in Supplementary Figure S4.

| Database | TF_ID | Class | MEME-ChIP<br>E-value | Overlap | offset | p-value | Tomtom<br>E-value | q-value |
| --- | --- | --- | --- | --- | --- | --- | --- | --- |
| chen2008 | **SMAD1** | **SMAD/NF-1 DNA-binding** | 8.4e-036 | **14** | **2** | **3.6e-03** | **4.3e-02** | 8.6e-02 |
|  | SOX2 | HMG domain |  | 13 | 2 | 1.8e-02 | 2.2e-01 | 2.2e-01 |
|  | OCT4 | Homeo domain |  | 13 | 2 | 2.9e-02 | 3.5e-01 | 2.2e-01 |
|  | NANOG | Homeo domain |  | 14 | 2 | 3.7e-02 | 4.4e-01 | 2.2e-01 |
| HOCOMOCO v11<br>(core) | **FOXJ3** | **Fork head/winged helix** |  | **13** | **-1** | **4.8e-06** | **1.7e-03** | **3.4e-03** |
|  | SRY | HMG domain |  | 9 | -4 | 3.9e-04 | 1.4e-01 | 1.4e-01 |
|  | AIRE | SAND domain |  | 14 | 0 | 1.5e-03 | 5.3e-01 | 2.0e-01 |
|  | FOXQ1 | Fork head/winged helix factors |  | 12 | -2 | 1.9e-03 | 6.8e-01 | 2.0e-01 |
|  | FOXJ2 | Fork head/winged helix factors |  | 10 | -1 | 2.0e-03 | 7.3e-01 | 2.0e-01 |
|  | NANOG | Homeo domain |  | 14 | 2 | 2.2e-03 | 7.7e-01 | 2.0e-01 |
|  | FOXL2 | Fork head/winged helix factors |  | 11 | -2 | 2.4e-03 | 8.7e-01 | 2.0e-01 |
|  | MEF2C | MADS box factors |  | 13 | 2 | 2.8e-03 | 9.9e-01 | 2.0e-01 |
| JASPAR 2022<br>(core) | **ZNF384** | **C2H2 zinc finger** |  | **12** | **-1** | **4.3e-06** | **8.4e-03** | **2.4e-03** |
|  | ONECUT1 | Homeo domain |  | 13 | -1 | 3.1e-04 | 6.0e-01 | 6.3e-02 |
|  | ONECUT3 | Homeo domain |  | 13 | -1 | 4.9e-04 | 9.6e-01 | 7.9e-02 |
| SwissRegulon | **ONECUT1** | **Homeo domain** |  | **12** | **-2** | **9.5e-05** | **1.8e-02** | **3.5e-02** |
|  | **BPTF** | **Bromodomain PHD zinc** |  | **14** | **0** | **2.5e-04** | **4.8e-02** | **4.6e-02** |
|  | IKZF2 | C2H2 zinc finger |  | 11 | 0 | 5.7e-04 | 1.1e-01 | 5.3e-02 |
|  | CDX1 | Homeo domain |  | 14 | 3 | 6.4e-04 | 1.2e-01 | 5.3e-02 |
|  | CUX2 | Homeo domain |  | 14 | 0 | 7.2e-04 | 1.4e-01 | 5.3e-02 |
|  | FOXI1 | Fork head/winged helix |  | 12 | -2 | 1.2e-03 | 2.2e-01 | 5.4e-02 |
|  | FOXO1 | Fork head/winged helix |  | 13 | -1 | 1.2e-03 | 2.2e-01 | 5.4e-02 |
|  | FOXD3 | Fork head/winged helix |  | 12 | -2 | 1.7e-03 | 3.3e-01 | 6.3e-02 |
|  | EVI1 | C2H2 zinc finger |  | 14 | 0 | 2.2e-03 | 4.1e-01 | 7.2e-02 |
|  | HMGA1 | A.T hook |  | 13 | -1 | 3.2e-03 | 6.1e-01 | 9.0e-02 |
|  | FOXQ1 | Fork head/winged helix |  | 11 | -2 | 4.4e-03 | 8.3e-01 | 1.1e-01 |

The MEME-ChIP analysis also revealed a similarity to the OCT4 binding site (Table 3), although this was not significant after correction for multiple testing. OCT4 (encoded by *POU5f1*) regulates itself and RNAPII-specific TFs, including NANOG and SOX2, and it plays an important role in maintaining chromatin structure in mESCs (Ding et al., 2012). Notably, OCT4, NANOG, SOX2 and AIRE are key regulators of mESC pluripotency and differentiation (Table 3), and the genes encoding these TFs are included in TG3 (Table 2 and Supplementary Table S1). Indeed, DIDO3 was shown to transcriptionally regulate *POU5f1*/OCT4 in mESCs (Fütterer et al., 2021). Collectively, these findings suggest the existence of a feedback loop between protein subunits in large RNAPII-associated complexes and their encoding genes that possess a DIDO3-binding site.

The possibility that DIDO3 might engage in non-specific DNA interactions was evaluated by analyzing the crystallographic structures available (Supplementary Figure S4B), performing a structural comparison between DIDO3-PHD in complex with H3K4me3 peptide (PDB ID: 4L7X) and ARID5-PHD in a ternary complex with H3K4me3 peptide and AT-rich DNA (PDB ID: 6LQF). Although DNA-binding specificity in ARID5 is conferred by its ARID domain, its neighboring PHD domain can form non-specific contacts with DNA. Interestingly, the positively charged DIDO3 residues R302 and R306 occupy positions equivalent to K712 and K716 in ARID5, which directly contact DNA, an emerging PHD-DNA interaction reviewed recently (Gaurav & Kutateladze, 2023).

### DIDO3 is associated with regulatory R-loops

R-loops are three-stranded nucleic acid structures consisting of an RNA:DNA hybrid and the displaced single-stranded DNA, classified as ‘unscheduled’ R-loops and scheduled, ‘regulatory’ R-loops. Regulatory R-loops are implicated in chromatin accessibility and transcriptional regulation, both by activating and silencing gene expression (Niehrs & Luke, 2020). Recent studies reveal that chromatin states influence R-loop dynamics, whereby H3K36me3-marked active genes exhibit R-loop resistance while H3K27ac-enriched enhancers show R-loop susceptibility (Yan et al., 2020). In DIDO3ΔE16 ESCs there is enhanced nuclear R-loop accumulation (Fütterer et al., 2021), although the underlying mechanism remains unclear. Several genes classified here in TG1 were considered R-loop regulators in human and mouse according to the R-loopBase (*Carm1*, *Dhx16*, *Smarcc1*, *Trim28*, *Wdr33*), or in TG3 (*CENPA*, *Cenpv*, *Ddx5*, *Dhx30*, *DDX39B*, *DNMT3B*, *Ep400*, *Ezh2*, *Hdac1*, *PABPC1*, *PAF1*, *PRPF4*, *Prpf6*, *Set*, *Setx*, *SMARCA5*, *Smc3*, *SRSF1*, *SRSF2*, *Suz12*, *Sox2* and *Upf1*: Lin et al., 2022).

The amino acid sequences of DIDO3 from different mammalian organisms were compared and a RGG/RG rich domain found was also present in EWSR1, a multifunctional RNA-binding protein that plays a critical role in R-loop regulation, genome stability and maintaining centromere identity (Figure 5: Kitagawa et al., 2023; Lay et al., 2024). In EWSR1, both the RGG/RG domain and RNA-recognition motif (RRM) confer specificity to R-loop binding. Moreover, the conserved DIDO3-CT RGG/RG rich domain includes an amino acid sequence of ancient *Ornithorhynchus anatinus* (platypus), annotated as a “single-stranded DNA binding protein” region (SSDP, a.a. 1926-2200: Figure 5). Although DIDO3 does not have a RRM motif, the RGG/RG rich domain could contribute to R-loop recognition. Moreover, there is evidence that various confirmed interactors of DIDO3 (CTCF, DHX9, HNRNPU, SFPQ) act as R-loop regulators (Petti et al., 2019; Lin et al., 2022; Ferrando et al., 2026) and independent studies demonstrate that R-loops, together with CTCF, are boundaries of cohesin-mediated DNA looping (Zhang et al., 2022). Together, these observations are consistent with a possible association of DIDO3 with 3D-chromatin structure and transcriptional regulation.

**Figure 5.**
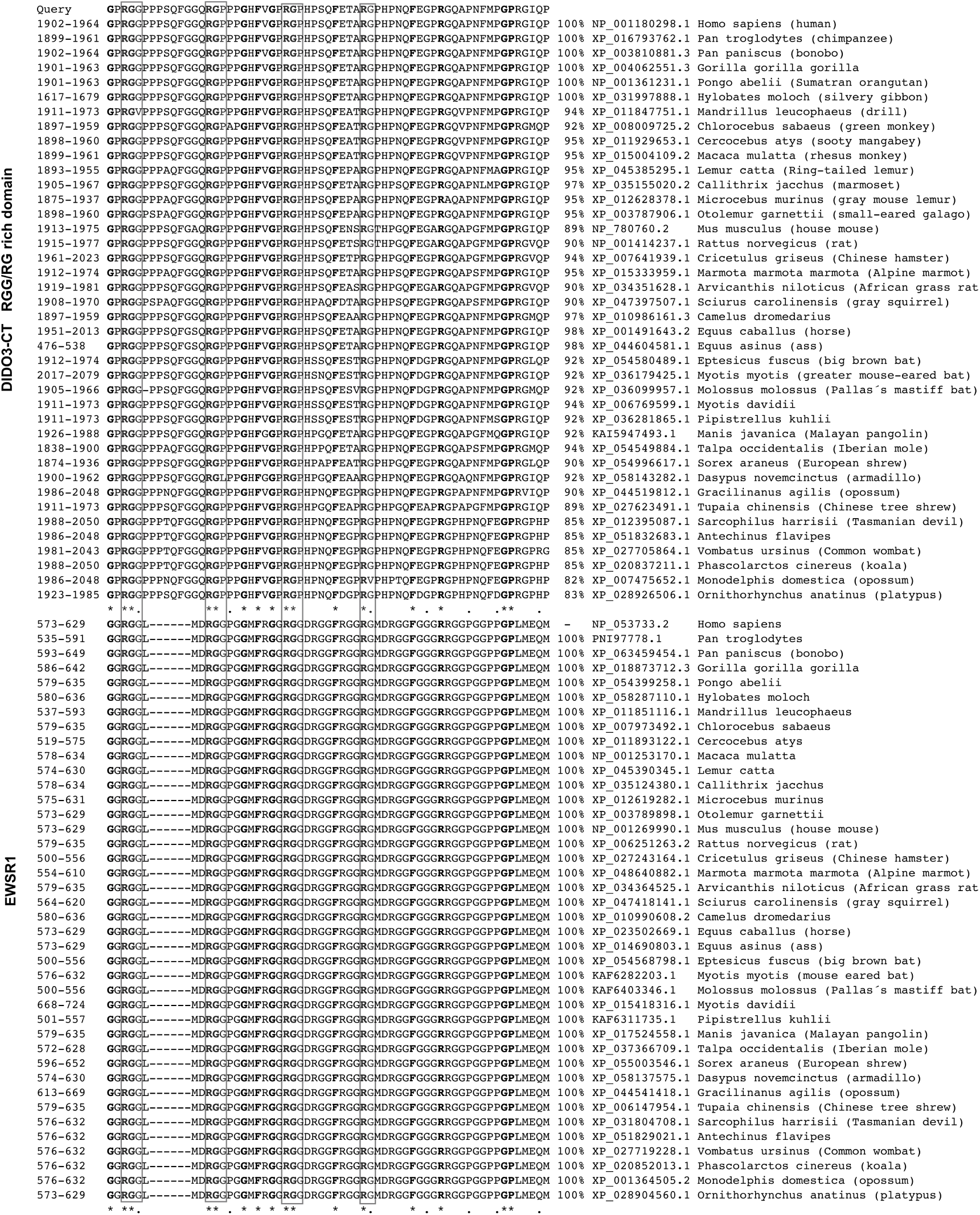
DIDO3 CT contains a conserved RGG/RG rich domain in mammals. Comparison of the amino acid sequences of mammalian DIDO3 CT domains and EWSR1. Asterisks indicate identical residues and the dots mark highly conserved positions. The numbers correspond to the first and last residues of each domain in the mature proteins. Residues conserved between the two families are shown in bold and the conserved RGG/RG motifs are boxed. The RefSeq identifiers, organism names and sequence identity is also shown.

### DIDO3 and RNAPII spatially coexist during the cell cycle and are immediately adjacent to the CREST signal

The localization of DIDO3 and RNAPII in MEFs (where DIDO3 is the predominant isoform: Fütterer et al., 2005) was assessed by confocal fluorescence microscopy, ensuring precise characterization with an antibody against the DIDO NT (recognizing all DIDO isoforms), a CT specific antibody for DIDO3 and an anti-HA antibody fibroblasts expressing HA-DIDO3. The autoimmune CREST serum, which recognizes a range of centromeric and kinetochore proteins (Moroi et al., 1980), was also used as a structural marker to delimit the condensed centromeric chromatin. The analysis of prometaphase cells revealed that DIDO3 and RNAPII coincide, yet only partially with the CREST marker (Figure 6A, left panel). Quantitative line-scan analysis showed that the bulk of the DIDO and RNAPII signals are situated between distances of 0.0 and 0.8 μm, effectively placing them outside the primary CREST peak concentrated between 0.9 and 1.1 μm.

**Figure 6.**
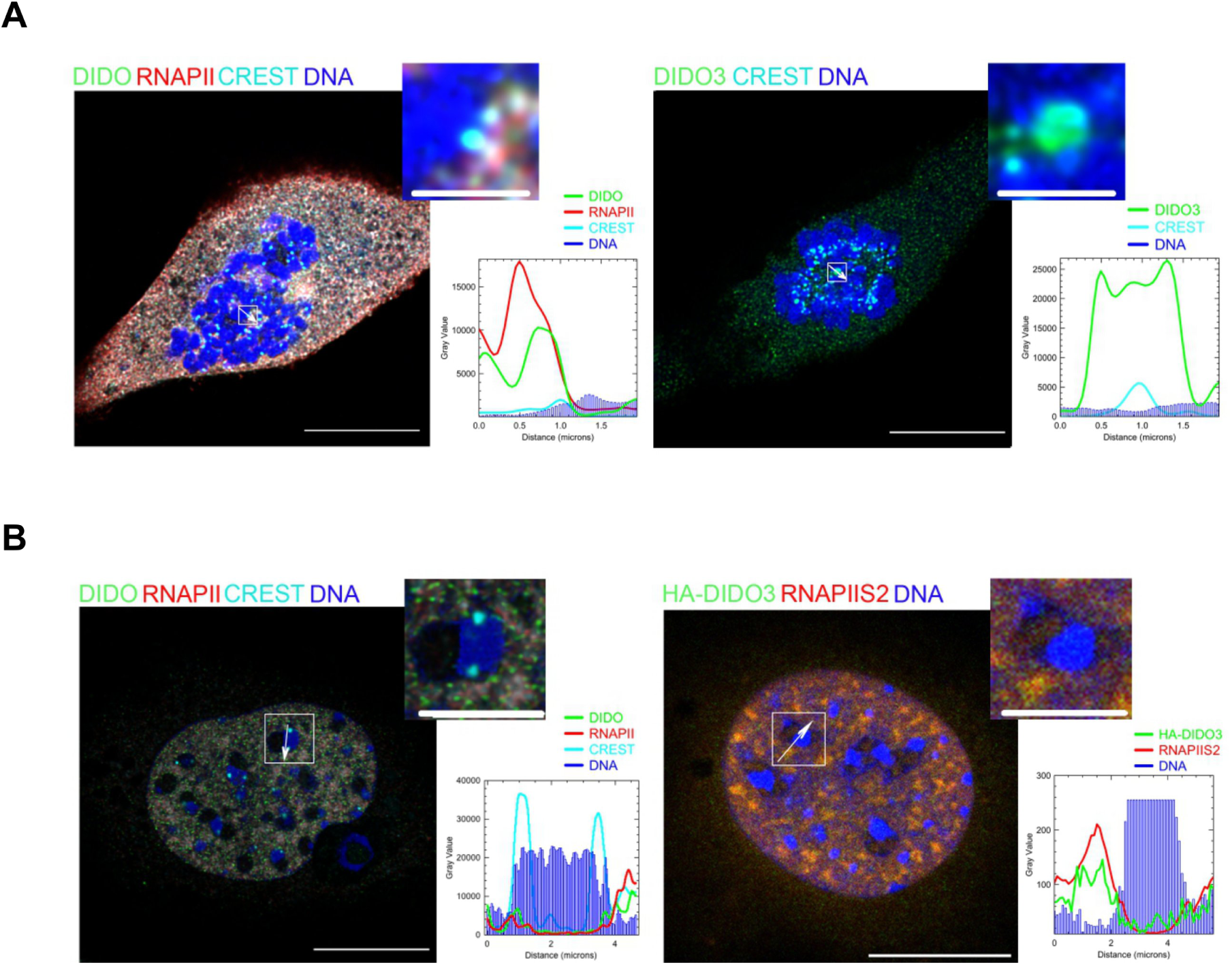
DIDO3 and RNAPII co-localize throughout the cell cycle, immediately adjacent to the CREST signal. Representative single stacks illustrating the distribution of DIDO (green), RNAPII (red) and CREST (cyan) in mitotic cells during prometaphase (A) and at interphase (B). In (A), the left panel shows DIDO detected with an anti-DIDO antibody and RNAPII at kinetochores (CREST), while the right panel shows DIDO3 detection with an anti-DIDO3 antibody and CREST. In (B), wild-type MEFs are labeled for DIDO (green), RNAPII (red) and CREST (cyan) in the left panel, or HA-DIDO3 and active RNAPIIS2 in the right panel. Scale bars: 15 μm for the main images; insets A 2 μm and B 5 μm. The plots illustrate the intensities of DIDO, RNAPII and CREST in the regions indicated by arrows in the boxed areas, with the 0 coordinate on the X-axis corresponding to the starting point of the arrow.

This spatial arrangement is also maintained during interphase, consistent with the findings that presence of RNAPII transcriptional activity in mouse fibroblasts at this stage (Chan et al., 2012). DIDO3 and RNAPII co-localized in a domain immediately adjacent to the condensed centromeric DNA marked by CREST (Figure 6B, left panel). In fibroblasts expressing HA-DIDO3, a robust overlap with RNAPIIS2ph was detected (a marker of transcriptional elongation) spanning a region between 0.5 and 2.2 μm (Figure 6B, right panel). While RNAPIIS2ph has been associated with an active kinetochore (Chan et al., 2012), it is important to note that this modification is a general hallmark of active RNAPII during gene body transcription and it is not exclusive to kinetochore structures.

In conclusion, these data demonstrate that DIDO3 and active RNAPII are spatially associated within a broad domain adjacent to the condensed centromeric chromatin. This specific localization of DIDO3 and active RNAPII in a broad domain adjacent to the condensed centromeric core (CREST) suggests several key functional roles, such as the regulation of centromeric transcription. RNAPII transcription at centromeric, and largely pericentromeric regions, is primarily dictated by the epigenetic state of chromatin rather than the DNA sequence.

### Similar epigenetic patterns also suggest co-regulation of DIDO3 and chromatin-associated complexes in human cells

As DIDO3ΔE16-induced alterations in key chromatin regulatory mechanisms were identified in mice, the presence of similar regulatory mechanisms in human cells was examined using the Epicorr v1.20 methodology in a genome-wide search for correlated chromatin marks over the *Dido1* locus and other regions (see Methods). The *Dido1* locus has a 10 kb variable epigenetic pattern across samples in part of the locus (∼61,545 kb in chromosome 20: Figure 7, top) and scanning all 1 kb windows within this fragment against all 1 kb windows of the 24 human chromosomes, 167 pairs of genomic regions with a correlation score >0.7 were identified (a complete list of these regions is available at http://csbg.cnb.csic.es/pazos/DIDO1_epigen/). Among these correlated regions, enhancers, promoters and coding regions associated to the human *ARID1A*, *BCL11A*, *CENPA*, *CENPE*, *CENPF*, *HJURP* and *CTBP2* genes were found (Figure 7 and Supplementary Table S3). *ARID1A* and *BCL11A* are components of the SWI/SNF complex, whereas *CTBP2* functions as an accessory protein of PRC2.1. CENPA serves as a centromeric marker and in mammals, it associates with the HJURP chaperone that mediates its deposition and maintenance within chromatin (Dunleavy et al., 2009; Foltz et al., 2009). In HeLa cells, active RNAPII can physically associate with CENPA-containing chromatin during early G1 (Quénet & Dalal, 2014). *CENPE* and *CENPF* are centromere-associated proteins essential for proper microtubule-kinetochore function. Additional genes in TG1 to TG3 display correlation scores between 0.6 and 0.7, such as *JARID2*, *BIRC5*, *CENPM*, *WAPL* and *WDR82* (Supplementary Table S3). These genes participate in diverse chromatin- and centromere-related processes, including association with PRC2.2 (*JARID2*), recruitment of the Chromosomal Passenger Complex to centromeres (*BIRC5*), contribution to the CENPA nucleosome-associated complex involved in kinetochore assembly (*CENPM*), regulation of cohesin loading onto chromatin (*WAPL*), or participation in the Set1C/COMPASS complex by binding to the RNAPII C-terminal domain (*WDR82*). Interestingly, the homologous mouse genes *ARID1A*, *CENPA*, *CTBP2*, *HJURP, JARID2* and *WAPL* had DIDO3 binding sites in WT ESCs (Table 2). Indeed, *Dido1* generates the same three protein isoforms by alternative splicing in mouse and human cells (Fütterer et al., 2005), and mouse and human DIDO3 sequences are 75% identical. These results suggest a possible co-regulatory role for DIDO3, HJURP/CENP-A chaperone complex, SWI/SNF, Set1C/COMPASS and PRC2 in both murine and human cells.

**Figure 7.**
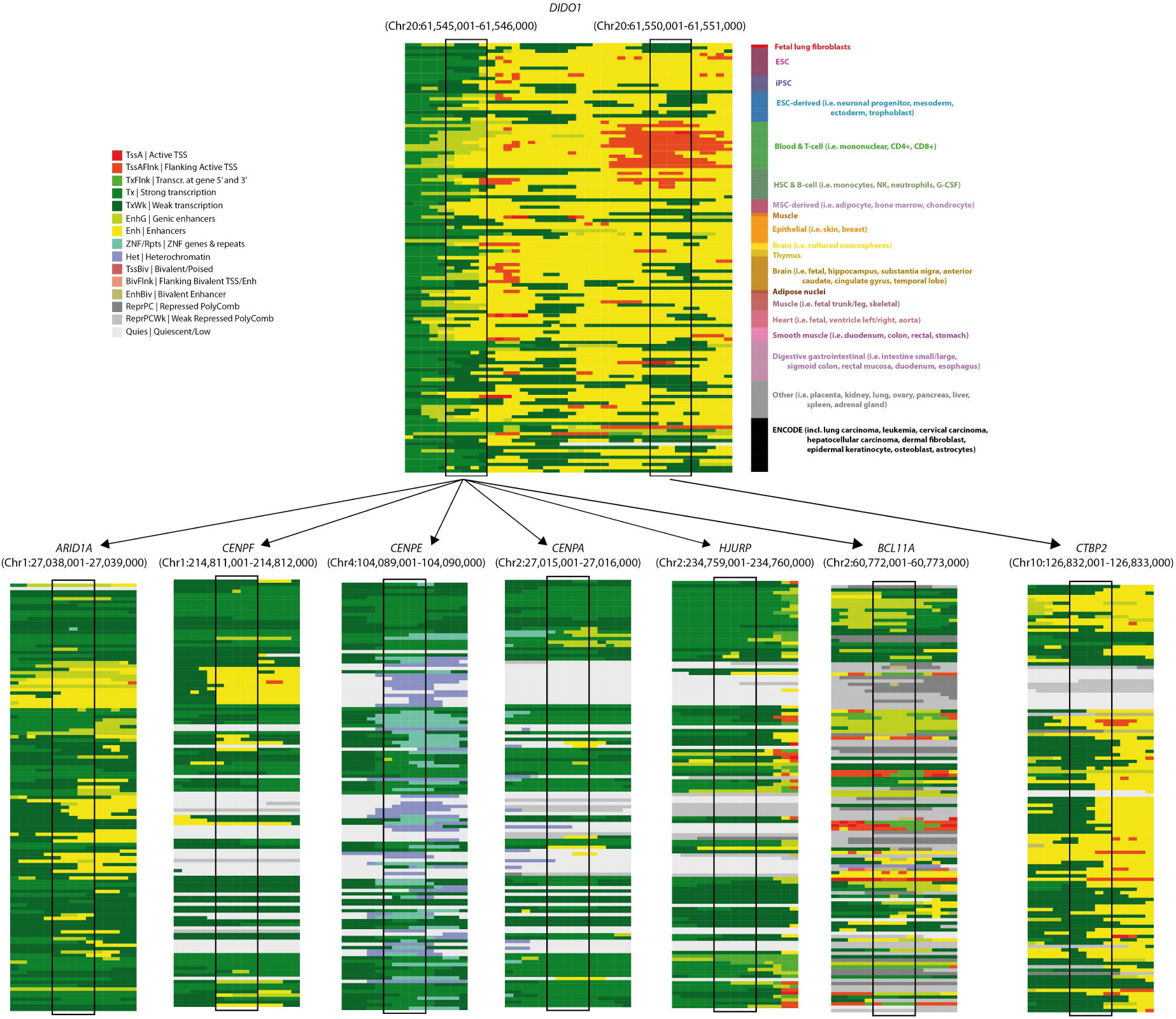
Genomic regions with informative epigenetic patterns. The epigenetic state of each 200 bp portion (small boxes) is indicated using the color code indicated in the figure. The two *Dido1* regions discussed (1,000 bp windows) are highlighted in large boxes, surrounded by their epigenomic neighbourhood. The names of the epigenetic sample groups are in the top right corner, colored using the tissue-based color scheme of the Roadmap Epigenomics Consortium.

To further analyze these 167 regions with an epigenetic pattern correlated with that in the *Dido1* locus, the genes matching these regions were retrieved and a functional enrichment analysis was carried out (Supplementary File 3). Two main categories were found in the list of significant terms: i) cell adhesion and signaling (orange, Supplementary File 3); and ii) cell division and chromosome segregation (green, Supplementary File 3). These two categories match the centromere-localized DNA breaks (Guerrero et al., 2010) and cell differentiation defects in the DIDO3 mutant (Fütterer et al., 2017; 2021), and the results suggest a possible association between DIDO3 and centromere transcription in human cells.

### Recurrent alterations accumulate at the DIDO3 splice-site and in the CT domain of distinct cancer types

Deregulation of chromatin structure is associated with cancer development (Feng & Pauklin, 2020; Jerković et al., 2020) and almost every tumor acquires somatic mutations in TFs, chromatin regulators or the DNA regulatory elements critical for transcriptional control (Doane & Elemento, 2022). *Dido1* gene alterations were recently reported in small cohorts of cancer patients (Table 4) although *Dido1* is not classified as a potential cancer driver. The distribution of mutations in the DIDO3 amino acid sequence was studied in at least two different samples and of the 6 truncating mutations in *Dido1*, four resided in the DIDO3 CT domain after the PHD, TFSII_M and SPOC domains, comparable to the DIDO3ΔE16 mutant (p.G1745Afs*119, p.P1908Lfs*111, p.P2034Rfs*240 and the splice site p.X1181_splice in Figure 8A).

**Figure 8.**
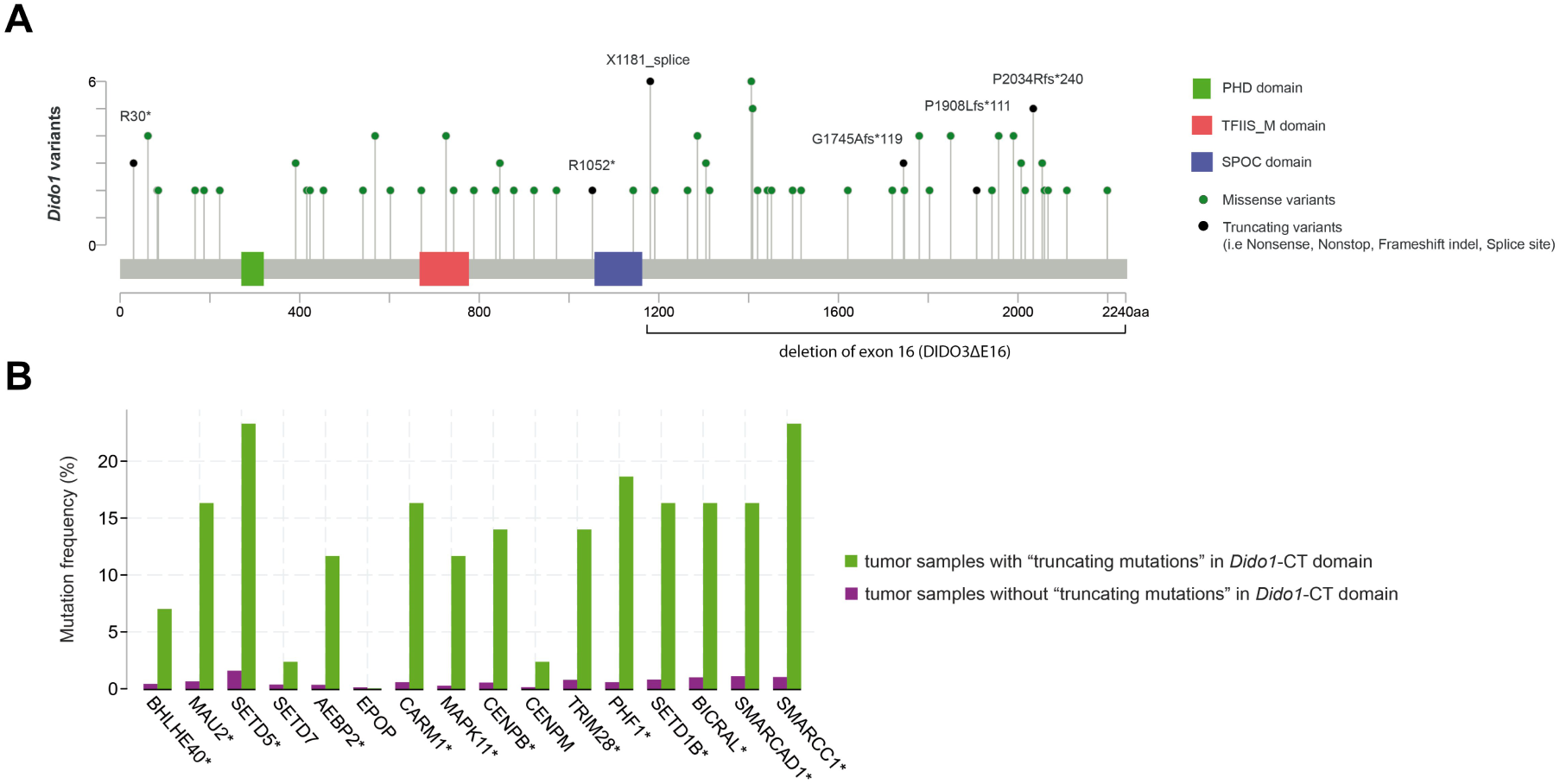
*Dido1* genetic alterations in the TCGA-PanCancer dataset. (A) Lolliplots showing the distribution of coding variants (missense, nonsense and frameshift) over the DIDO3 functional domains. (B) Mutation frequency of DIDO3 primary and secondary targets in tumor samples with “truncating mutations” in the DIDO3 CT domain: * indicates statistical significance (Fisher’s exact test, P <0.001).

**Table 4.**
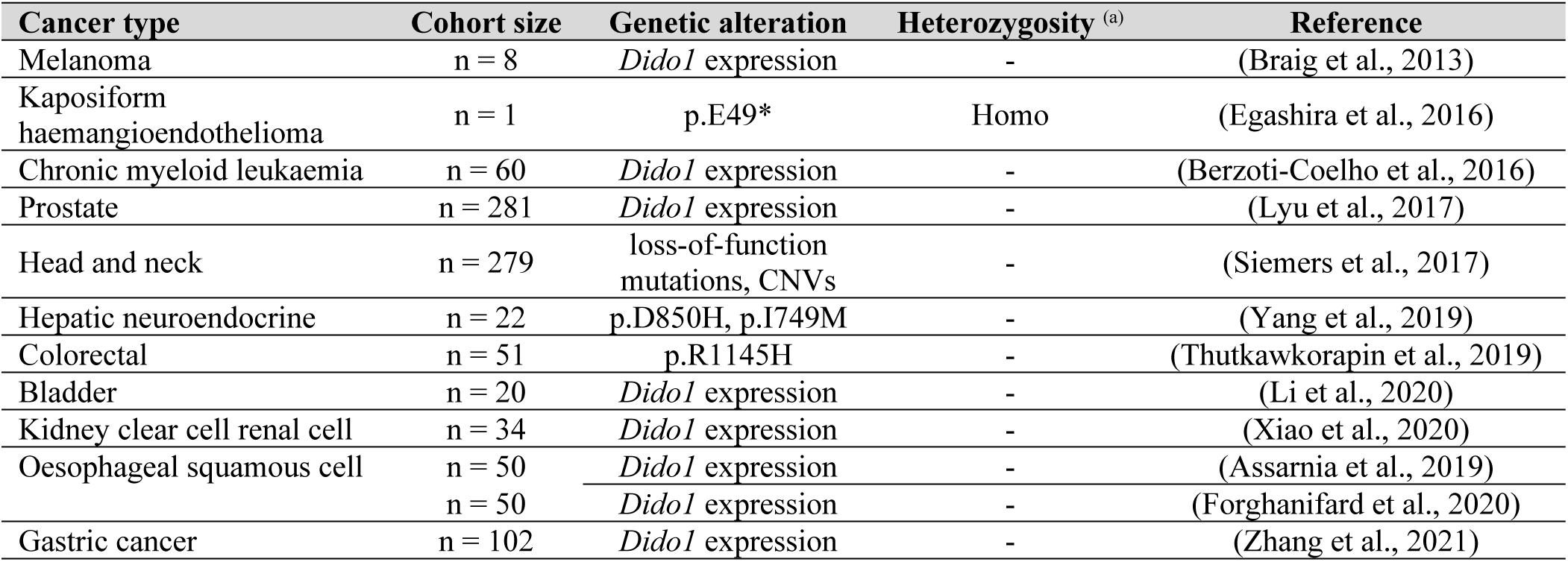
Cancer types with *Dido1* genetic alterations reported in the literature. ^(a)^ The two alleles present at each *Dido1* locus contain the genetic variant. A Hyphen (-) indicates that information about heterozygosity is not provided.

Changes to the expression of genes encoding components of the CTCF-cohesin, PRC2, SWI/SNF, Set1C/COMPASS and centromere-associated protein network were evident with the DIDO3ΔE16 mutant. Each of these macromolecular complexes coordinate their activities to regulate chromatin structure and gene expression (Rittenhouse et al., 2025; Jerković et al., 2020; Müller & Almouzni, 2017). Based on the TCGA-PanCancer dataset, tumor samples with mutations truncating the DIDO3 CT domain (black dots, Figure 6A) also accumulate mutations in genes like: *BHLHE40* and *MAU2* related to CTCF-cohesin complex; *AEBP2*, *EPOP* and *PHF1* related to PRC2; *BICRAL*, *SMARCAD1* and *SMARCC1* related to SWI/SNF; *SETD1B* related to Set1C/COMPASS; *CENPB* and *CENPM* components of the centromere protein network; and the chromatin regulators *SETD5*, *SETD7* and TRIM28. These genes are the human homologues of DIDO3 primary and secondary target genes in mouse ESCs (Table 1). Except for *CENPM*, *EPOP* and *SETD7*, the remaining 13 genes accumulated significantly more mutations in tumor samples with truncating mutations in the DIDO3 CT domain (Fisher’s exact test, P <0.001, according to cBioPortal: Figure 8B). These observations again point to a possible association between DIDO3 and large macromolecular complexes that may be involved in chromatin structure and gene expression dynamics in mice and humans.

## DISCUSSION

Gene expression and repression are tightly regulated processes essential for cell differentiation and development. These processes are governed by a network of chromatin modifiers, TFs, RNAPII, ‘regulatory’ R-loops and architectural proteins (including CTCF and cohesin complexes) that together help organize the three-dimensional structure of the genome. Aberrant activity of chromatin modifiers can drive epigenetic silencing or inappropriate gene activation, a hallmark of many diseases like cancer (Stojchevski et al., 2025). Oncogenic TFs may hijack normal developmental pathways to promote tumor growth (Marchetto et al., 2020) and mutations in CTCF binding domains or cohesin subunits can alter the higher-order organization of the genome, resulting in misexpression of critical genes (Song et al., 2017). In addition, RNAPII dysregulation can provoke pauses, elongation or termination that it turn may disrupt transcriptional programs and contribute to malignant transformation (Bradner et al., 2017). Despite advances in understanding these factors, the precise mechanisms by which chromatin modifiers, transcriptional machinery, regulatory R-loops and genome architecture interact to shape gene expression programs are still not fully understood, particularly in the context of cancer.

Our RNA-seq data revealed that genes encoding multifunctional, multi-subunit protein complexes involved in epigenetic regulation, chromatin remodeling (such as SWI/SNF and NuRD) and histone modification (including the Polycomb and MLL-family histone methyltransferase complexes) were overrepresented among the up- and downregulated genes in DIDO3ΔE16 mutants. Many of these complexes interact with RNAPII and play a pivotal role at the interface of transcriptional regulation and three-dimensional genome architecture (Batsché et al., 2006; Ui et al., 2016). The dual role of these complexes in chromatin dynamics and transcriptional control suggests that they may be important modulators of cancer related gene expression (Bracken et al., 2019).

RNAPII transcription not only occurs at protein-coding and non-coding genes but also, pervasively across intergenic regions (Gajos et al., 2021). The DIDO3 protein illustrates this regulatory complexity: its TFIIS_M domain binds directly to RNAPII (Matveyeva et al., 2025); and as seen here, its PHD domain binds to histone H3-tail peptides (Gatchalian et al., 2013), co-localizing with H3K4me3 and H3K36me3, while its CT domain interacts with the architectural CTCF protein. Indeed, the DIDO3ΔE16 truncation affects RNA splicing and transcription termination in mESCs and MEFs (Fütterer et al., 2021; Mora Gallardo et al., 2021; Mora Gallardo et al., 2019). Notably, CTCF itself can regulate alternative splicing, further contributing to the complexity of transcriptome remodeling in cancer (Alharbi et al., 2021).

Together, these findings indicate that chromatin modifiers, transcriptional machinery and genome architecture components co-localize, such that perturbations in any of these systems could influence programs of gene expression.

### Novel (CTCF and H3K36me3) and confirmed (RNAPII, H3K27ac and H3K4me3) DIDO3 targets

The ChIP-seq analysis indicates an overlap between the chromatin binding sites of DIDO3, RNAPII, H3K36me3, H3K27ac, H3K4me3 and CTCF in mESCs. The co-localization of DIDO3 and H3K4me3 appears to be particularly strong, consistent with experimental data regarding PHD domain specificity (Gatchalian et al., 2013). An overlap was also found between DIDO3 and the PRC2 repressor complex, although the analysis could not determine whether this reflects direct physical interactions or simply co-occupancy of shared genomic regions. Notably, the overlap between H3K4me3 and PRC2 is unexpected (Schmitges et al., 2011) but may be due to the frequent presence of both at promoter regions that might harbor both activating and repressive chromatin features (Cookis et al., 2025).

To further study co-localization and distinguish specific patterns, the genome was divided into promoter and intergenic regions. DIDO3 did not co-localize with any transcriptional repression marks or proteins in intergenic regions, suggesting that its association with PRC2 at promoters may be coincidental, as may also be the case for H3K4me3. However, strong co-localization between DIDO3 and H3K36me3 was evident in intergenic regions where the latter did not overlap with any other marks or proteins tested, consistent with a potential association between DIDO3 and H3K36me3. Indeed, the DIDO3 PHD domain is very similar to other PHD domains known to specifically recognize H3K36me3 (Shi et al., 2007). These observations were validated by confocal microscopy and SPRi, collectively supporting the possible co-localization of DIDO3 with H3K36me3.

PHD fingers exhibit considerable diversity in their ability to recognize a wide range of PTMs and unmodified histone tails (Li et al., 2012), including di- or tri-methylated K4, trimethylated K9, acetylated K14 and trimethylated K36, as well as H4 modifications like acetylation (Bortoluzzi et al., 2017). The presence of multiple positively charged lysine and arginine residues at the surface of PHD fingers suggests that these domains may also interact with DNA and indeed, mutations of lysine residues in the α1 region of the BAZ1A PHD finger disrupt its DNA-binding capacity (Oppikofer et al., 2017).

### A feedback loop between RNAPII transcription and DIDO3 targets

The interplay between RNAPII pausing and chromatin structure regulation has been studied (Francette et al., 2021) and here we suggest that DIDO3 may co-localize with factors involved in RNAPII pausing and the regulation of chromatin structure. Based on the predicted association of DIDO3 with H3K36me3 and the interaction of its CT-domain with CTCF, DIDO3 may be associated with genomic regions involved in RNA splicing and transcription termination (Fütterer et al., 2021; Mora Gallardo et al., 2021; Mora Gallardo et al., 2019). Although the DIDO3 PHD domain is known to bind primarily to H3K4me3 (Gatchalian et al., 2013), H3K36me3 itself has been implicated in cryptic transcription, alternative pre-mRNA splicing and 3D chromatin organization (Fang et al., 2021; Leung et al., 2019). Furthermore, CTCF mediates the RNAPII elongation rate in a methylation-dependent manner (Alharbi et al., 2021), RNAPII pausing (Chernukhin et al., 2007; Gajos et al., 2021; Luan et al., 2021) and it influences RNA splicing events (Alharbi et al., 2021; Shukla et al., 2011), although the precise mechanisms remain to be elucidated. The DIDO3 CT domain not only alters interactions with the SFPQ splicing factor (Mora Gallardo et al., 2019), CTCF (this study) and the DHX9 helicase (Fütterer et al., 2021), but it may also co-localize with other proteins essential for RNAPII processing (e.g., HDAC1, CDK12, PAF1). ChIP-seq analysis revealed that the genes encoding several of these proteins contain DIDO3-binding sites, including: the largest subunit of RNAPII (RPB1/Polr2a), the splicing factor Sfpq, the helicase Dhx9, the H3K27ac-specific histone deacetylase Hdac1, the cyclin-dependent kinase Cdk12, and the RNAPII-associated factor Paf1. While RNA-seq did not show significant expression changes in DIDO3ΔE16 cells, the proximity of DIDO3 to these genes suggests that further studies will be needed to determine the functional consequences of DIDO3 binding at these loci.

### Role of DIDO3 in 3D chromatin structure

Aggregate peak analysis (APA) identified 3D co-localization patterns in the linear genomic sequence similar to those observed through overlapping ChIP-seq peaks. DIDO3 is enriched at sites with indicators of transcriptional activity, like H3K4me3, H3K27ac, H3K36me3, RNAPII, and active promoters and enhancers. To a lesser extent, DIDO3 co-localizes with repressive Polycomb complexes and H3K27me3, and a less direct interaction between DIDO3 and CTCF was observed than reported previously when a clear overlap of their binding sites was noted. Hence, the association between DIDO3 and CTCF may be indirect, potentially due to DIDO3 binding at promoters located near but not precisely at CTCF-CTCF loop anchors. Notably, the presence of CTCF at anchor points was negatively correlated with DIDO3, which was more frequently positioned toward the interior of the loop (25-100 kb from the anchor point). A similar spatial displacement was observed for RNAPII and histone marks associated with active transcription, an effect most pronounced for DIDO3. DIDO3 was clearly enriched at TAD borders, consistent with CTCF protein and transcriptional activity. DIDO3 appeared to be enriched in A compartments (generally associated with active chromatin) as opposed to B compartments with inactive chromatin. Moreover, the genes with DIDO3 marks upregulated in DIDO3ΔE16 cells were more likely to be located in B compartments, whereas downregulated genes were more frequently found in the active A compartment. Notably, those among the former with weaker expression were less likely to be located in B compartments.

### Ideas and Speculation

DIDO3 functions as an accessory factor for RNAPII and it acts as a chromatin reader, yet the mechanisms through which DIDO3 (and the DIDO3ΔE16 mutant) modulate transcription and chromatin architecture are only partially understood. Integrating ChIP-seq, RNA-seq and Hi-C analyses in mESCs and MEFs with previous studies indicates DIDO3 may act through several mechanisms. In terms of RNAPII pausing, the 3D organization of chromatin can regulate the elongation rate of RNAPII, thereby influencing alternative splicing. DIDO3 may participate in transcription-coupled methylation/deacetylation, potentially enabling the rapid repression of induced genes once their expression is no longer required. DIDO3 binding sites were identified in the *Nsd1* gene encoding a H3K36me2-specific methyltransferase and in *Hdac1*, a H3K27ac-specific deacetylase. In addition, several histone methyltransferases were identified as TG1 or TG2 targets, including *Setd7* (H3K4me1), *Setd1b* (H3K4me2/3) and *Setd5* (H3K36me3). The elongation rate of RNAPII can also be affected by the interaction between DIDO3 and CTCF (Alharbi et al., 2021). The participation of DIDO3 in transcription-coupled methylation/deacetylation may contribute to restoring a repressive chromatin state, which could be involved in silencing cryptic promoter-like sequences within gene bodies and repressing antisense transcription in coding regions (Chantalat et al., 2011). This hypothesis is supported by the co-localization of DIDO3 with H3K4me3 and H3K36me3, and its potential interaction with HDAC1. Both H3K36me3 and histone deacetylase complexes can work together to modulate expression in complex feedback circuits (Fang et al., 2021; Keogh et al., 2005). After the DIDO3 PHD-domain recognizes H3K36me3, its CT domain and that of RPB1 may recruit HDAC1 and lead to deacetylation of H3K27 (see Supplementary Figure S1 in Pons et al., 2014). This mechanism could prevent transcription initiation from cryptic promoter-like sequences within the body of genes. Indeed, the gene encoding PAF1, involved in modulating RNAPII elongation rates, was seen to also contain a DIDO3 binding site in WT ESCs (TG3 group).

PRC2 is a chromatin-modifying complex that enforces transcriptional silencing through epigenetic regulation and its two major functional variants, PRC2.1 and PRC2.2, share a conserved catalytic core but incorporate distinct accessory subunits. PRC2.1 predominantly maintains repression once it has been established, whereas PRC2.2 is more closely associated with *de novo* silencing of active loci. Here, the core PRC2 components EED, EZH2 and SUZ12 were identified as TG3 targets, along with accessory factors from both assemblies. PRC2.2-associated proteins like AEBP2 and JARID2, and the PRC2.1-associated proteins like CTBP2, ELOB, ELOC, EPOP, PHF1 and MTF2, were considered TG1 or TG3 targets. Moreover, *Phf1* (an H3K36me3 reader) was downregulated, while *Aebp2* and *Epop* were upregulated in DIDO3ΔE16 mutant cells.

The ATP-dependent SWI/SNF complexes (bBAF, GBAF, nBAF, npBAF) enhance promoter accessibility and facilitate RNAPII transcription. By antagonizing PRC1 and PRC2, SWI/SNF helps maintain the regulatory balance between RNAPII activity and Polycomb repression, and disruption of this equilibrium contributes to oncogenesis (Bracken et al., 2019). Both shared and complex-specific SWI/SNF components were identified as TG1 (*Gltscr1l*/*Bicral*, *Smarcad1*, *Smarcc1*) and TG3 (*Arid1a*, *Bcl7b*, *Smarca5-ps, Smarce1*, *Phf10*) targets. In DIDO3ΔE16 mutant cells, *Gltscr1l*/*Bicral* and *Smarcad1* were upregulated, whereas *Smarcc1* was downregulated. In addition, the Epicorr analysis of various human cell types reveals a pattern of chromatin marks at the *DIDO1* locus and the genomic regions of *ARID1A*.

Finally, the mESC ChIP-seq and RNA-seq data identified centromere-associated proteins as TG2 (*Cenpb*, *Cenpm*) and TG3 (*Cenpa*, *Cenph*, *Cenpv*) targets, with *Cenpb* and *Cenpm* upregulated in DIDO3ΔE16 mutant cells. Previous studies with DIDO3 mutants reported double-strand DNA breaks near centromeres, which were linked to mitotic spindle defects (Guerrero et al., 2010). In addition, other genes involved in the centromere-associated protein network (*Hjurp*, *Mis18bp1*, *Dnmt3b*) also have DIDO3 binding sites. Indeed, the Epicorr analysis of various human cell types reveals a correlated pattern of chromatin marks at the *DIDO1* locus, and at the *CENPA*, *CENPE*, *CENPF* and *HJURP* genomic regions.

## CONCLUSION

In summary, our findings suggest that DIDO3 associates with RNAPII, and it co-localizes with the H3K4me3 and H3K36me3 chromatin marks that are involved in transcription and regulating chromatin structure. The integration of experimental data suggests that DIDO3 may regulate chromatin structure through its co-localization with promoter-enhancer loops and its association with CTCF, cohesin loading complexes, SWI/SNF and Polycomb complexes. As such, DIDO3 may play a role in pathways involved in cell differentiation and it may be relevant for cancer development. Nevertheless, further experimental studies will be required to elucidate the precise mechanisms by which DIDO3 influences RNAPII processivity and chromatin structure.

## MATERIALS AND METHODS

A detailed description of the software, databases and computational data processing methods used for ChIP-seq, RNA-seq and Hi-C experiments, as well as the *de novo* motif discovery, protein-interaction analysis and analysis of epigenetically correlated regions is provided in Appendices 1 and 2. Protein codes and functional annotations were obtained from UniProtKB (UniProt Consortium, 2018), Pfam (Mistry et al., 2021) and PROSITE (Hulo et al., 2006). Differential gene expression analysis was performed as described elsewhere (Fütterer et al., 2021). Protein co-localization was assessed in single-cell regions of fluorescence images using a custom Java-based script compatible with the ImageJ and Fiji platforms. The script used for the data analysis is available at https://github.com/QuantitativeImageAnalysisUnitCNB/XYProteinColocalization. For each cell, a specific region of interest (ROI) was defined in deconvoluted images and the Pearson’s correlation coefficient was calculated for the volume. Nuclei were detected in the DAPI channel using ImageJ, and foci in the red and green channels were counted using a user-defined threshold above background (detailed methods are provided in Appendix 1).

### Experimental procedures

#### qRT-PCR

Total RNA was extracted using the guanidinium isothiocynate method and the “HOT FIREPol EvaGreen qPCR Mix Plus (5x)” was used for PCR reactions, which were run on an Applied Biosystems QuantStudio^TM^ 5 Real-Time PCR System, with QuantStudio^TM^ Design & Analysis v1.5.0 software. The ThermoFisher Cloud Relative Quantification tool was used to analyze the results applying the Comparative Ct Method. Detailed procedures and primer sequences are provided in Appendix 2.

#### Cell culture and cell transfections

MEFs were cultured on coverslips in high glucose Dulbecco′s Modified Eagle′s Medium (DMEM) supplemented with L-glutamine and 5% fetal bovine serum (FBS). Mammalian expression vectors, including pCAGG-HA-tagged DIDO3 (Fütterer et al., 2012) and DIDO3_1-528_ with either the WT PHD or the C285A/H290A mutated PHD domain (Gatchalian et al., 2013), were transiently transfected into MEFs using the Lipofectamine 2000 reagent and according to the manufacturer’s protocol. For western blotting or co-immunoprecipitation experiments, ESCs were cultured as described elsewhere (Gatchalian et al., 2013). Further details are provided in Appendix 2.

#### Immunofluorescence

For histone labeling, cells were fixed for 10 minutes with ice-cold methanol, while for other immunofluorescence labeling, the samples were fixed in 3.7% formaldehyde for 15 minutes at room temperature (RmT). In both cases, the cells were permeabilized with 0.1% Triton X-100 in PBS for 5 minutes, and blocked with 3% BSA and 0.05% Tween 20 in PBS for 20 minutes prior to antibody incubation. The list of antibodies used is provided in Appendix 2. Images were acquired using a Leica STELLARIS 5 confocal multispectral system equipped with four laser lines, three Power HyD S spectral detectors, and an integrated software module for real-time super-resolution image detection and processing.

## DATA AVAILABILITY

The complete list of correlated genomic regions identified using Epicorr v1.20 is available at http://csbg.cnb.csic.es/pazos/DIDO1_epigen/ and on Zenodo (https://zenodo.org/records/22093186). Confocal images are available from the BioStudies database (https://www.ebi.ac.uk/biostudies/) under accession numbers S-BIAD1354 and S-BIAD1390. Unless otherwise noted, all other datasets used in this study are available from the Gene Expression Omnibus (GEO). ChIP-seq data are available under the following accession numbers: GSE85029 (HA-DIDO3), GSM2645517 (Rpb1), GSM2645511 (H3K36me3), GSM2645495 (H3K4me3), GSM2645513 (H3K27ac), GSM2645497 (H3K27me3), GSM2645515 (CTCF), GSE120393 (PRC2), and GSM2645509 (Ring1B). RNA-seq data for DIDO3 and DIDO3ΔE16 in mouse embryonic stem cells (mESCs) are available under accession number GSE152346; corresponding data in mouse embryonic fibroblasts (MEFs) are available from the NCBI Sequence Read Archive under accession numbers SRP150516 and PRJNA476070. Hi-C data are available under accession numbers GSE99530 (mESC), GSE121087, and GSE115983 (MEF).

## SUPPLEMENTARY DATA

Supplementary Data is available at eLife Online

## AUTHOR DETAILS

### Tirso Pons

**Contribution:** Conceptualization, Methodology, Data curation, Investigation, Formal analysis, Writing – original draft preparation, Writing—review and editing, Supervision.

**Competing interests:** No competing interests to declare.

### François Serra

**Contribution:** Conceptualization, Methodology, Software, Investigation, Formal analysis, Writing – original draft preparation, Writing—review and editing.

**Competing interests:** No competing interests to declare.

### Agnes Fütterer

**Contribution:** Investigation, Formal analysis, Visualization, Resources, Writing—review and editing.

**Competing interests:** No competing interests to declare.

### Karel H. M. van Wely

**Contribution:** Formal analysis, Resources, Writing—review and editing.

**Competing interests:** No competing interests to declare.

### Florencio Pazos

**Contribution:** Methodology, Software, Investigation, Formal analysis, Writing—review and editing, Funding acquisition.

**Competing interests:** No competing interests to declare.

### Cristina Pacios-Bras

**Contribution:** Investigation, Formal analysis, Visualization, Writing—review and editing.

**Competing interests:** No competing interests to declare.

### Ana Cayuela López

**Contribution:** Methodology, Software, Investigation, Formal analysis, Writing—review and editing.

**Competing interests:** No competing interests to declare.

### Carlos Oscar S. Sorzano

**Contribution:** Formal analysis, Resources, Writing—review and editing.

**Competing interests:** No competing interests to declare.

### Alfonso Valencia

**Contribution:** Resources, Writing—review and editing. **Competing interests:** No competing interests to declare.

### Carlos Martínez-A

**Contribution:** Conceptualization, Writing—review and editing, Project administration, Supervision, Funding acquisition.

**Competing interests:** No competing interests to declare.

## ACKNOWLEDGMENTS

The authors thank all members of the Stem Cells and Immunity research group for early sharing of information and for their feedback on our manuscript. We thank Luis Almonacid for assistance with the qRT-PCR experiments and the Advanced Light Microscopy Unit for providing us the dataset of confocal images. We also thank Catherine Mark and Dr Mark Sefton for editorial assistance.

## FUNDING

This work was supported by the Spanish State Research Agency (AEI/10.13039/501100011033) through grants SAF2016-75456-R and PID2019-110574RB-I00 (to C.M.A. and K.v.W.), PID2019-108096RB-C22 (to F.P.), and the “Severo Ochoa” Programme for Centres of Excellence in R&D (CEX2023-001386-S). Additional funding was provided by the Comunidad de Madrid (P2022/BMD-7321/MITIC-CM) and the Intramural Project (PIE 201920E017 to C.M.A.).

## APPENDIX 1

### Software

This study utilized a range of bioinformatics software tools to analyze genomic and epigenomic data. For ChIP-seq data analysis and binding site annotation, we used ChIPseeker (Yu et al., 2015), clusterProfiler (Yu et al., 2012) and BEDTools (Quinlan & Hall, 2010). Motif discovery and enrichment analysis of transcription factor (TF) binding sites were performed with MEME-ChIP (Machanick & Bailey, 2011). PyRanges (Stovner & Sætrom, 2020) was employed for efficient comparison of genomic intervals, while NetworkX (Hagberg et al., 2008) was used for graphical representation. TADbit (Serra et al., 2017) enabled the analysis of chromatin three-dimensional interaction matrices, including the identification of topologically associated domains (TADs) and A/B compartments from Hi-C experiments. TADs are regions of the genome that interact more frequently with themselves than with other regions (Dixon et al., 2012). A/B compartments are broad domains identified in Hi-C data: A compartments correspond to transcriptionally active chromatin, which is generally gene-rich and associated with open chromatin marks; B compartments correspond to transcriptionally inactive chromatin, which is gene-poor and associated with repressive epigenetic marks (Lieberman-Aiden et al., 2009). Metascape (Zhou et al., 2019) provided comprehensive gene list annotation, functional enrichment and interactome analysis. Epicorr v1.20 (Pazos et al., 2018) was used to identify regions of the human genome epigenetically correlated with the Dido1 locus. Functional enrichment analysis was performed with the topGO R package (bioconductor.org/packages/topGO/: Alexa et al., 2006) and RStudio (www.rstudio.org) was used to run R/Bioconductor packages, including org.Mm.eg.db, TxDb.Mmusculus.UCSC.mm10.knownGene, ggplot2 and edgeR.

### Databases

To compare well-characterized sequence-specific binding sites of mouse TFs, we used the following databases: HOCOMOCO v11 (Kulakovskiy et al., 2018), JASPAR 2022 (Castro-Mondragon et al., 2022), SwissRegulon (Pachkov et al., 2013) and the Chen2008 collection of embryonic stem cell (ESC) TFs (Chen et al., 2008) All databases were accessed via the MEME Suite 5.4.1 tool (http://meme-suite.org: Bailey et al., 2015). Cancer-associated variants were obtained from the Pan-Cancer TCGA dataset using cBioPortal (www.cbioportal.org: Cerami et al., 2012). Protein codes and functional annotations were sourced from UniProtKB (www.uniprot.org: UniProt Consortium, 2018), Pfam (pfam.xfam.org/: Mistry et al., 2021) and PROSITE (prosite.expasy.org: Hulo et al., 2006).

### Data processing

#### ChIP–seq data analysis

The genomic coordinates for HA-DIDO3 binding (GSE85029) were converted to the UCSC mouse genome build mm10 using LiftOver (genome.ucsc.edu/cgi-bin/hgLiftOver). BED files were then imported into RStudio and annotated using the R/Bioconductor package ChIPseeker (Yu et al., 2015), with the promoter region defined as -1 kb to 200 bp relative to the transcription start site (TSS). Additional peak annotations were performed using the Bioconductor packages org.Mm.eg.db and TxDb.Mmusculus.UCSC.mm10.knownGene. The significance of overlap between ChIP-seq datasets was assessed using the enrichPeakOverlap function in ChIPseeker. This function compares the observed overlap between peaks to a null distribution generated by 10,000 random permutations (nShuffle=10000) to assess if the overlap is greater than expected by chance. Overlapping genomic sites were identified using the intersectBed function in BEDTools (Quinlan & Hall, 2010) and PyRanges (Stovner & Sætrom, 2020) was used for further comparison of genomic intervals.

#### RNA-seq and differential gene expression analysis

RNA-seq data for DIDO3 and DIDO3ΔE16 in mouse ESCs (GSE152346) and MEFs (SRP150516 and PRJNA476070) were processed as previously described (Fütterer et al., 2021). Briefly, paired-end reads were aligned to the UCSC mouse genome build mm10 using BWA-MEM 0.7.15 (http://bio-bwa.sourceforge.net) with default settings. The resulting alignments were converted to BAM format and de-duplicated using Picard tools 2.9.0 (http://broadinstitute.github.io/picard/). Transcript abundance was quantified with StringTie 1.3.3 (Pertea et al., 2015) and expression levels were calculated as transcripts per million (TPM). Only transcripts with TPM >0 in all samples were retained for downstream analysis. Data normalization and differential expression analysis were performed using the R package edgeR (Robinson et al., 2010). Genes were considered differentially expressed if they exhibited an absolute log2 fold change of at least 0.7 and a false-discovery rate (FDR) below 0.05.

#### Protein-protein interaction, gene ontology and pathway analysis

The complete list of genes with HA-DIDO3 binding sites detected in ChIP-seq, as well as those with gene expression alterations in RNA-seq data, was submitted to Metascape (Zhou et al., 2019) using the multi-gene-list and meta-analysis option. Metascape integrates protein interaction data from BioGRID (thebiogrid.org) and generates a protein-protein interaction (PPI) network, identifying modules (sub-networks of highly connected proteins) using the MCODE clustering algorithm. Gene ontology (Ashburner et al., 2000) and pathway enrichment analyses were also performed with Metascape. For enrichment analysis, we used the core set of default ontologies provided by Metascape, including Gene Ontology (GO) processes, KEGG pathways, Reactome gene sets, canonical pathways and CORUM protein complexes. The hypergeometric test (Zar & Zar, 1999) and Benjamini-Hochberg p-value correction algorithm (Benjamini et al., 1995) were used by Metascape to identify ontology terms significantly enriched in the input gene list.

#### Hi-C data processing

All the Hi-C data collected from GEO was first converted to FASTQ files using parallel-fastq-dump. Raw reads were processed with TADbit (Serra et al., 2017) for quality control, read mapping, interaction detection, interaction filtering and matrix normalization. Valid interactions were then used to generate genome-wide interaction maps at 100 kb, 50 kb, 20 kb and 5 kb resolution. These maps were used to segment the genome into A/B compartments and TADs, as well as to perform meta-analyses. A/B compartments were calculated using normalized and decay-corrected matrices, with a principal component analysis (PCA) performed on chromosome-wide Pearson correlation matrices following the previously described method (Lieberman-Aiden et al., 2009) implemented in TADbit (Serra et al., 2017). TADs were identified at 50 kb resolution using the TAD border detection method in TADbit (Serra et al., 2017). Chromatin loops were called using the Mustache software (Roayaei Ardakany et al., 2020) at 10 kb resolution with default parameters, except that the sparsity threshold was lowered to 0.8 and the p-value threshold increased to 0.5. Aggregate peak analysis (APA) was performed by extracting small submatrices (21 x 21 cells at 5 kb resolution) from the genomic Hi-C matrix at the intersection of specific pairs of genomic coordinates (Rao et al., 2014). These submatrices, corresponding to all input pairs of genomic coordinates, were averaged into a single APA matrix representing the average interaction pattern between the two lists of input coordinates. To further analyze these interactions, we transformed the Cartesian coordinates of the APA matrix into a measure of distance from the center by calculating the Manhattan distance (the sum of the absolute distances along the X- and Y-axes) from the center cell (see Figure 2B for an example between DIDO3 ChIP-seq peaks and H3K36me3 ChIP-seq peaks). Based on this distance, measured in the number of 5 kb bins, interaction cells were classified into two categories: those close to the center (≤5 bins) and those farther away (>5 bins). We then compared the interaction strengths in these two categories using a two-sample t-test to assess whether there was significant enrichment of interactions at the center. A significant enrichment at the center of the APA indicates a higher frequency of interactions between the proteins or marks represented by the input coordinate lists, suggesting their co-localization in the nucleus.

#### Computational analysis of epigenetically correlated regions

To search for human genome regions epigenetically correlated with the *Dido1* locus, we used the epigenomic datasets compiled by the Roadmap Epigenomics Consortium (Kundaje et al., 2015). We downloaded the 127 “consolidated epigenomes”, for which the chromatin states are given in a 15-state vocabulary for all 200 bp (base-pair) windows (egg2.wustl.edu/roadmap/web_portal/chr_state_learning.html). The 127 epigenomes include different tissues, ESCs, cancer cell lines and developmental stages. For these epigenomes, detailed information on various epigenetic marks and features is provided at bp resolution, such as histone acetylation, DNA methylation, DNase accessibility, or RNA-seq expression. For each 200 bp genomic region, the ChromHMM method (Ernst & Kellis, 2017) is used to integrate all these experimental epigenetic proxies into a single “epigenetic state” with 15 possible values: active TSS (TssA), flanking active TSS (TssAFlnk), transcription at gene 5’ and 3’ (TxFlnk), strong transcription (Tx), weak transcription (TxWk), gene enhancers (EnhG), enhancers (Enh), Zinc Finger genes & repeats (ZNF/Rpts), heterochromatin (Het), bivalent/poised TSS (TssBiv), flanking bivalent TSS/enhancer (BivFlnk), bivalent enhancer (EnhBiv), repressed PolyComb (ReprPC), weak repressed PolyComb (ReprPCWk), and quiescent/low (Quies: Ernst & Kellis, 2017; Kundaje et al., 2015). The BED files with these chromatin states were downloaded for the 127 epigenomes. For lower resolution analyses using 1 kb windows, each window was assigned the most frequent epigenetic state among its constituent 200 bp segments if that state was present in at least 80% of the segments. If no single state met this criterion, the window was assigned an *undefined* state. The entire human epigenome was scanned using 1 kb non-overlapping windows. Each window’s epigenetic profile was defined as a vector of length 127, with each element representing the epigenetic state (one of 15 possible states) of that window in one of the 127 epigenomic samples. To assess the similarity between the epigenetic profiles of two genomic windows (*Gi* and *Gj*), we calculated the “mutual information” (MI), an entropy-based measure that quantifies the amount of shared information between their corresponding vectors, as follows:

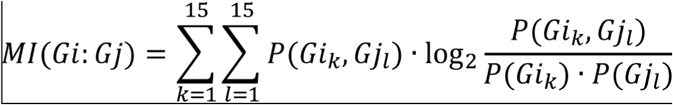

where the sums run for the 15 possible epigenetic states. *P(G_ik_)* and *P(G_jl_)* are the frequencies of epigenetic states *k* and *l* in windows *Gi* and *Gj*, respectively, and *P(G_ik_, G_jl_)* is the frequency of state *k* in window *Gi* matching state *l* in window *Gj* in the same samples. The *undefined* state is ignored for this calculation, and the terms of the sums where either *P(G_ik_)* or *P(G_jl_)* equal 0 are skipped. Thus, the mutual information parameter attains a high value when the epigenetic profiles of two windows are very similar. *<u>Fluorescence Image Analysis</u>*. A custom designed Java-based script was developed to operate within the ImageJ and Fiji ecosystem (github.com/QuantitativeImageAnalysisUnitCNB/XYProteinColocalization). This workflow implements pixel-by-pixel intensity correlation methods in batch mode across two different channels, allowing for the automatic selection of a specific region of interest (ROI) within the cell or alternatively, the entire cell, as predefined by the user. The workflow extracts each series from “.LIF” files and converts them into multichannel z-stack TIFF formats for downstream analysis. The “splitting multi-channel images” command is used to separate each z-stack volume into its red, green and blue channels, after which individual slices are extracted as independent images. To isolate ROIs (e.g. kinetochore) or whole cells for co-localization analysis, the “Auto-Threshold” plugin (Huang et al., 1995) is used to binarize images with the “Default” method (Ridler & Calvard, 1978). This is followed by the “Create Selection” command, which is used to count, measure and extract features of each ROI. ROIs that meet the filtering conditions are selected for further analysis. If a DAPI signal is present, only ROIs whose centroid falls within a DAPI-stained area are considered relevant, otherwise all the ROIs are included. The “Combine” command from the “ROI Manager” is then used to merge ROIs slice by slice using the union operator. Co-localization between X protein and Y protein is analyzed after background correction to account for spatial variations. At this stage, mean intensity values for each protein, along with co-localization indicators, are computed.

To analyze co-localization, Pearson’s and Manders’ coefficients were used to quantify the degree of overlap between fluorescent signals in confocal images acquired under identical conditions. Pearson’s coefficient assesses the linear correlation between the pixel intensities of X and Y proteins, with values ranging from 1 (perfect positive correlation), through 0 (no correlation), to -1 (perfect negative correlation).

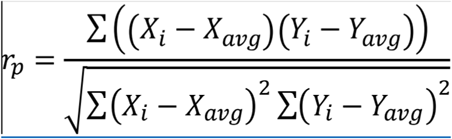

with 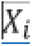 and 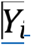 being the pixel values with the pixel index and 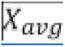 and 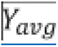 the averages of the 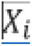 and 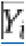 protein signals respectively, and the summations with index 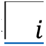 over all the image pixels.

Unlike the Pearson’s correlation coefficient, Manders’ overlap coefficients do not subtract the mean signal and therefore, they always yield values between 0 (no overlap) and 1 (complete overlap). These coefficients measure the fraction of total fluorescence from one protein that overlaps with the fluorescence of the other protein: M1 quantifies the proportion of X protein intensity overlapping with that of the Y protein, while M2 quantifies the proportion of Y protein intensity overlapping with the X protein.

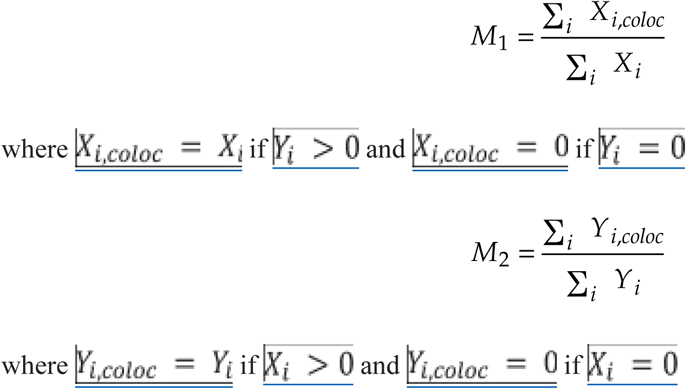

Finally, the p-value from the two-sample Kolmogorov-Smirnov test (with statistical significance indicated by *p* < 0.05: Massey, 1951) is computed to determine whether the fluorescence signals of the X and Y protein share the same spatial distribution. This test provides additional information beyond the co-localization analysis, as X and Y proteins may not co-localize but could still exhibit spatial dependence. Specifically, the test compares the distribution of pixel-by-pixel intensities (after subtracting the mean intensity for each protein) to the distribution of normalized pixel-by-pixel intensities of the X protein and the flipped version of the Y protein, as described above.

## APPENDIX 2

### Experimental procedures

#### qRT-PCR

Total RNA was extracted using the guanidinium isothiocynate method (TRIzol reagent, Invitrogen, Carlsbad, CA: Chomzynski & Sacchi, 1987) and approximately 1 μm was reverse transcribed with the High Capacity cDNA Reverse Transcription Kit (ThermoFisher-Applied Biosystems). The primer sequences used are: Aebp2-forward GCAGACCACATTCGCTCCATAC and Aebp2-reverse GCTGTAGCCAACTCTGACTGGT; Epop-forward CTTGACTGCTTCCCCTGTCC and Epop-reverse GTCCTCCCATCTGCCACTTC; GAPDH-forward AGAAACCCTGGACCACCCA and GAPDH-reverse CTCCCTCACAATTTCCATCCC; β-Actin-forward GGCTCCTAGCACCATGAAGA and β-Actin-reverse CCACCGATCCACACAGAGTA. All the PCRs were performed using “HOT FIREPol EvaGreen qPCR Mix Plus (ROX), 5x” (Solis Biodyne), the PCR reactions were run on an Applied Biosystems QuantStudio^TM^ 5 Real-Time PCR System (384-well), with QuantStudio^TM^ Design & Analysis v1.5.0 software, and the ThermoFisher Cloud Relative Quantification tool was used to analyze the results using the Comparative Ct Method (ΔΔCt: Livak & Schmittgen, 2001).

#### Cell culture and cell transfections

For immunofluorescence analysis, mouse embryonic fibroblasts (MEFs) were cultured on cover slips in high glucose Dulbecco′s Modified Eagle′s Medium (DMEM, Biowest, Cat No L0101) supplemented with L-glutamine (ApexBio, Cat No A8461) and 5% fetal bovine serum (FBS: Sigma Aldrich, Cat No F7524). The HA-DIDO3 (Fütterer et al., 2012), HA-NT and HA-NT with PHD mutation (Gatchalian et al., 2013) and HA-CT (Fütterer et al., 2017) plasmids were transiently transfected into MEFs using the Lipofectamine 2000 transfection reagent (ThermoFischer Sci, Cat No 11668019) according to the manufacturer’s protocol, generating stable ESC transfectants and selecting them with puromycin. For western blot or co-immunoprecipitation studies, ESCs were cultured as described elsewhere (Gatchalian et al., 2013) and to induce differentiation, these ESCs were cultured in MEF medium without LIF in low-adhesion plates. Lysates were probed with anti-histone antibodies using the histone extraction kit (ab113476, Abcam) and also with the H3K27 methylation sampler Kit (53312, Cell Signaling). In all cases, samples were separated by 4%–15% SDS-PAGE depending on protein size, transferred to nitrocellulose membranes (Bio-Rad) and probed with the antibodies indicated. ECL solution (PerkinElmer) was used to visualize proteins.

#### Immunofluorescence

For histone labelling, cells were fixed for 10 minutes in ice-cold methanol, while for other immunofluorescence labelling, samples were fixed in 3.7% formaldehyde (Sigma Aldrich, Cat No 47608) for 15 min at room temperature (RmT). In both cases, the cells were permeabilized with 0.1% Triton X-100 (Sigma Aldrich, Cat No 11332481001) in PBS for 5 minutes, and blocked in 3% BSA (Sigma Aldrich, Cat No A2153) and 0.05% Tween 20 (Sigma Aldrich, Cat No P1379) in PBS for 20 minutes before antibody incubation. Primary antibodies (Abs) were incubated for either 2 hours at RmT or for 16 hours at 4 °C, followed by three 5 minute washes with PBS. Secondary Abs were then incubated for 1-2 hours at RmT and washed as before, and the labelled coverslips were mounted in DAPI-containing ProLong Gold (Invitrogen, Cat No P36930). Images were acquired with a confocal multispectral system Leica STELLARIS 5, equipped with 4 laser lines (405, 488, 561 and 638 nm), 3 Power HyD S spectral detectors, and an integrated software module (Lightning) for real-time multidimensional super-resolution image detection and processing. These images are available in the BioStudies database (www.ebi.ac.uk/biostudies/) under accession numbers S-BIAD1354 and S-BIAD1390. The images in Figure 6 (panel A) were deconvoluted with Huygens, and ROIs and plots were generated with Image J software (Schneider et al., 2012) before they were assembled in Adobe Photoshop (www.adobe.com/products/photoshop.html). In Figure 6B, right panel, the acquisition channels were inverted with ImageJ for clarity. As such, HA labelled with the 546 secondary Ab is shown in green and RNAPIIS2 labelled with the 488 secondary Ab is shown in red. The primary Abs used were: Anti-HA.11 monoclonal Ab (clone 16B12, BioLegend) diluted 1:500; anti histone 3 trimethylated lysine specific Abs, such as anti-H3K4me3 rabbit monoclonal Ab (ab213224, Abcam) diluted 1:500, the anti-H3K36me3 rabbit monoclonal Ab (ab282572, Abcam) diluted 1:500, the anti-H3K27me3 rabbit polyclonal antiserum (9756, Cell Signaling) diluted 1:500 and the anti-H3K9me3 rabbit polyclonal antiserum (ab8898, Abcam) diluted 1:5,000; Anti-DIDO monoclonal Ab (Prieto et al., 2009) diluted 1:100, Anti-DIDO3 polyclonal antiserum (Prieto et al., 2009), Anti-RNAPII (N20, sc-899, Santa Cruz Biotechnology); Anti-RNAPIIS2ph (ab5095, Abcam), Anti-HA-Biotin, 3F10 (12158167001, Merck) diluted 1:100; CREST serum (del Mazo et al., 1987) diluted 1:5,000. The secondary Abs used were: Goat anti-mouse IgG (H+L) Alexa Fluor 488 (ThermoFisher Scientific, Cat No A-11029), diluted 1:400; Cy™3 AffiniPure goat anti-rabbit IgG (Jackson Immunoresearch, Cat No 111-165-008), diluted 1:200 and goat anti-human Cy5 (Jackson Immunoresearch, Cat No 109-165-098), diluted 1:800.

## Notes

https://www.ebi.ac.uk/biostudies/bioimages/studies/S-BIAD1354

https://www.ebi.ac.uk/biostudies/bioimages/studies/S-BIAD1390

https://github.com/acayuelalopez/XYProteinColocalization

https://zenodo.org/records/22093186

### Competing Interest Statement

The authors have declared no competing interest.

### Summary of Updates

The Data Availability section has been revised to clarify the origin of the datasets (newly generated vs. previously published). A public Zenodo URL has also been included for the complete set of correlated genomic regions identified via Epicorr v1.20; Author affiliations updated.

