## Supplementary material for "A genome-wide analysis identifies DIDO3 binding at active chromatin and topological boundaries in mouse stem cells": Supplementary_Information.pdf

**Supplementary Table S1. Primary and secondary target genes differentially expressed between DIDO3ΔE16 and wild-type ESCs, and genes with DIDO3 binding sites that may interact with DIDO3**

| ***Gene*** | ***Chr*** [(a)] | ***ChIP-seq*** [(b)] | ***RNA-seq*** [(c)] | ***Description*** [(d)] |
|---|---|---|---|---|
| **TG1 group** | | | | |
| *Jade1* | Chr3 (+) | #1708 to #1713 | $log_2FC$ = -5.91<br>FDR = 2.74e-04 | HBO1 complexes (histone H4 acetylation) |
| *Klf2* | Chr8 (+) | #2656, #2657 | $log_2FC$ = 0.7<br>FDR = 2.95e-03 | Transcription factor |
| *Mbd3* | Chr10 (-) | #226, #227, #228 | $log_2FC$ = 0.81<br>FDR = 1.02e-03 | NuRD complex (methyl-CpG binding, nucleosome remodeling, histone deacetylation, and DNA repair) |
| *Pus3* | Chr9 (+) | #2744, #2745, #2746 | $log_2FC$ = 3.94<br>FDR = 2.3e-09 | tRNA pseudouridine synthase |
| *Wdr33* | Chr18 (+) | #1300 | $log_2FC$ = 0.96<br>FDR = 6.47e-04 | mRNA 3'-end processing |
| **TG2 group** | | | | |
| *Foxj2* | Chr6 (+) | - | $log_2FC$ = -0.99<br>FDR = 4.85e-02 | Transcriptional coactivator |
| *Otx2* | Chr14 (-) | - | $log_2FC$ = -2.16<br>FDR = 2.76e-03 | Transcription factor (differentiation-associated) |
| *Suv420h1/Kmt5b* | Chr19 (+) | - | $log_2FC$ = -2.2<br>FDR = 9.52e-06 | Histone methyltransferase (H4K20me1/2) |
| **TG3 group** | | | | |
| *Cdk12* | Chr11 (+) | #521, #522, #523 | - | RNAPII CTD phosphorylation (transcription elongation, genomic stability, DNA repair) |
| *Cpsf4* | Chr5 (+) | #2187, #2188 | - | CPSF complex (pre-mRNA 3'-end formation) |
| *Cpsf6* | Chr10 (-) | #282 to #285 | - | CFIm complex (pre-mRNA 3'-end cleavage and polyadenylation) |
| *Ep400* | Chr5 (-) | #2124 | - | NuA4 complex (histone H2A and H4 acetylation) SWR1-like complex (removes H2A.Z/H2AZ1) |
| *Trrap* | Chr5 (+) | #2186 | - | NuA4 complex (histone H2A and H4 acetylation) SWR1-like complex (removes H2A.Z/H2AZ1) |
| *Esrrb* | Chr12 (+) | #659 to #663 | - | Transcription factor (self-renewal, pluripotency) |
| *Maz* | Chr7 (-) | #2537 | - | Transcription factor |
| *Npm1* | Chr11 (-) | #355, #356, #357 | - | Chaperonin for H3, H2B and H4, centrosome duplication, ribosome biogenesis |
| *Pabpc1* | Chr15 (-) | #924 | - | Binds the poly(A) tail of mRNA, regulates pre-mRNA splicing and mRNA stability |
| *Pabpn1* | Chr14 (+) | #854, #855 | - | 3'-end formation of mRNA precursors. Stimulates poly(A) polymerase |
| *Paf1* | Chr7 (+) | #2381 | - | PAF1 complex (histone H2B ubiquitination, H3K4 methylation). Modulates RNAPII elongation rates and ESC pluripotency |
| *Pnrc2* | Chr4 (-) | #1945, #1946, #1947 | - | Nonsense-mediated mRNA decay, nuclear receptor coactivator |
| *Smg5* | Chr3 (+) | #1744 | - | Nonsense-mediated mRNA decay |
| *Upf1* | Chr8 (-) | #2631 | - | Nonsense-mediated decay of mRNAs containing premature stop codons. RNA-dependent helicase |
| *Rpa2* | Chr4 (+) | #1925 | - | Core component of RNA polymerase I |
| *Sirt3* | Chr7 (-) | #2566, #2567 | - | $NAD^+$-dependent deacetylase. Involved in stabilizing heterochromatin and senescence |
| *Smad3* | Chr9 (-) | #2784 | - | Heteromeric SMAD protein complex. Transcription factor |
| *Suv39h1* | ChrX (-) | #2850 | - | Histone methyltransferase (H3K9me1), constitutive heterochromatin at pericentric and telomere regions |
| *Tet1* | Chr10 (-) | #194, #195, #196 | - | DNA demethylation. Chromatin regulation. Promotes histone H2B GlcNAcylation |
| *Zc3h12a* | Chr4 (-) | #1901 | - | Endoribonuclease (mRNA decay). Cleaves mRNA harbouring a stem-loop located in the 3'-UTR. |

[(a)] Gene position is represented by the chromosome (Chr) number and strand directions in parentheses as forward (+) or reverse (-). [(b)] Unique MACS2 identifiers correspond to the ChIP-seq peak from the HA-DIDO3 dataset (GSE85029). [(c)] Gene expression changes between DIDO3ΔE16 and wild-type embryonic stem cells, including log2 fold change ($log_2FC$) and adjusted p-values (false discovery rate, FDR) that were calculated from the RNA-seq data using edgeR. [(d)] Protein annotations from UniProtKB (www.uniprot.org). A hyphen (-) indicates that no ChIP-seq peak or gene expression change was detected for the corresponding entry.

**Supplementary Table S2. Intersection of DIDO3 binding sites with RNAPII, CTCF, Polycomb, and histone H3 modifications**

| PyRanges | | | | | | | |
|---|---|---|---|---|---|---|---|
| qSample[(a)] | tSample[(b)] | qLen[(c)] | tLen[(d)] | Overlap (%)[(e)(f)] | All[(g)] | Promoters[(g)] | Intergenic[(g)] |
| DIDO3 | RNAPII | 2 888 | 29 041 | 2 269 (78.6%) | +++ | +++ | +++ |
| DIDO3 | H3K36me3 | 2 888 | 33 970 | 2 002 (69.3%) | +++ | +++ | +++ |
| DIDO3 | H3K27ac | 2 888 | 29 284 | 1 036 (35.9%) | +++ | +++ | +++ |
| DIDO3 | H3K4me3 | 2 888 | 43 051 | 811 (28.1%) | +++ | +++ | +++ |
| DIDO3 | CTCF | 2 888 | 49 203 | 360 (12.5%) | +++ | +++ | +++ |
| DIDO3 | PRC2 | 2 888 | 23 646 | 242 (8.4%) | +++ | +++ | NS |
| DIDO3 | RING1B | 2 888 | 10 116 | 41 (1.4%) | NS | +++ | --- |
| DIDO3 | H3K27me3 | 2 888 | 19 812 | 22 (0.8%) | --- | NS | --- |
| RNAPII | H3K36me3 | 29 041 | 33 970 | 16 326 (56.2%) | +++ | +++ | --- |
| RNAPII | H3K27ac | 29 041 | 29 284 | 14 402 (49.6%) | +++ | +++ | +++ |
| RNAPII | H3K4me3 | 29 041 | 43 051 | 16 511 (56.9%) | +++ | +++ | +++ |
| RNAPII | CTCF | 29 041 | 49 203 | 7 174 (24.7%) | +++ | +++ | --- |
| RNAPII | PRC2 | 29 041 | 23 646 | 3 385 (11.7%) | +++ | +++ | --- |
| RNAPII | RING1B | 29 041 | 10 116 | 3 871 (13.3%) | +++ | +++ | --- |
| RNAPII | H3K27me3 | 29 041 | 19 812 | 2 310 (8.0%) | +++ | +++ | --- |
| H3K36me3 | H3K27ac | 33 970 | 29 284 | 8067(23.7%) | +++ | +++ | --- |
| H3K36me3 | H3K4me3 | 33 970 | 43 051 | 9 286 (27.3%) | +++ | +++ | --- |
| H3K36me3 | CTCF | 33 970 | 49 203 | 6 792 (20.0%) | +++ | +++ | --- |
| H3K36me3 | PRC2 | 33 970 | 23 646 | 1 557 (4.6%) | --- | +++ | --- |
| H3K36me3 | RING1B | 33 970 | 10 116 | 611 (1.8%) | --- | NS | --- |
| H3K36me3 | H3K27me3 | 33 970 | 19 812 | 449 (1.3%) | --- | --- | --- |
| H3K4me3 | H3K27ac | 43 051 | 29 284 | 18 893 (43.9%) | +++ | +++ | +++ |
| H3K4me3 | CTCF | 43 051 | 49 203 | 11 714 (27.2%) | +++ | +++ | +++ |
| H3K4me3 | PRC2 | 43 051 | 23 646 | 10 043 (23.3%) | +++ | +++ | +++ |
| H3K4me3 | RING1B | 43 051 | 10 116 | 10 236 (23.8%) | +++ | +++ | +++ |
| H3K4me3 | H3K27me3 | 43 051 | 19 812 | 9 754 (22.7%) | +++ | +++ | +++ |
| H3K27ac | CTCF | 29 284 | 49 203 | 6 848 (23.4%) | +++ | +++ | +++ |
| H3K27ac | PRC2 | 29 284 | 23 646 | 2 368 (8.1%) | +++ | +++ | --- |
| H3K27ac | RING1B | 29 284 | 10 116 | 2 636 (9.0%) | +++ | +++ | NS |
| H3K27ac | H3K27me3 | 29 284 | 19 812 | 1 121 (3.8%) | NS | +++ | --- |
| CTCF | PRC2 | 49 203 | 23 646 | 3 671 (7.5%) | +++ | +++ | +++ |
| CTCF | RING1B | 49 203 | 10 116 | 3 574 (7.3%) | +++ | +++ | +++ |
| CTCF | H3K27me3 | 49 203 | 19 812 | 4 279 (8.7%) | +++ | +++ | +++ |
| PRC2 | RING1B | 23 646 | 10 116 | 7 638 (32.3%) | +++ | +++ | +++ |
| PRC2 | H3K27me3 | 23 646 | 19 812 | 8 009 (33.9%) | +++ | +++ | +++ |
| RING1B | H3K27me3 | 10 116 | 19 812 | 7 142 (70.6%) | +++ | +++ | +++ |
| ChIPseeker | | | | | | | |
| qSample[(a)] | tSample[(b)] | qLen[(c)] | tLen[(d)] | N_OL[(e)] | N_OL (%qLen)[(f)] | p-value[(h)] | p-adjust[(i)] |
| HA-DIDO3 | RNAPII | 2 888 | 29 041 | 2 290 | 79.3% | 9.9e-05 | 1.6e-04 |
| HA-DIDO3 | H3K36me3 | 2 888 | 33 970 | 2 019 | 69.9% | 9.9e-05 | 1.6e-04 |
| HA-DIDO3 | H3K27ac | 2 888 | 29 284 | 1 050 | 36.4% | 9.9e-05 | 1.6e-04 |
| HA-DIDO3 | H3K4me3 | 2 888 | 43 051 | 829 | 28.7% | 9.9e-05 | 1.6e-04 |
| HA-DIDO3 | CTCF | 2 888 | 49 203 | 368 | 12.7% | 4.6e-03 | 6.9e-03 |
| HA-DIDO3 | PRC2 | 2 888 | 23 646 | 287 | 9.9% | 9.9e-05 | 1.6e-04 |
| HA-DIDO3 | RING1B | 2 888 | 10 116 | 41 | 1.4% | 0.2 | 0.2 |
| HA-DIDO3 | H3K27me3 | 2 888 | 19 812 | 22 | 0.8% | 0.9 | 0.9 |

[(a)] Query ChIP-seq sample. [(b)] Target ChIP-seq sample. [(c)] Number of query peaks. [(d)] Number of target peaks. [(e)] Number of overlapping peaks between query and target. [(f)] Percentage of overlapped peaks. [(g)] Genomic region analysed. [(h)] P-value calculated by ChIPseeker. [(i)] P-value correction (FDR) by the Benjamini-Hochberg method. Values were obtained using the ChIPseeker command *enrichPeakOverlap*, which assesses peak overlap through 10,000 random permutations of genomic locations (queryPeak=file1, targetPeak=file-list, TxDb= TxDb.Mmusculus.UCSC.mm10.knownGene, pAdjustMethod="BH", nShuffle=10000, chainFile=NULL, verbose=FALSE). Symbols indicate the degree of overlap: +++, more than expected (Bonferroni-corrected p<0.001); ++, more than expected (p<0.01); +, more than expected (p<0.05); ---, less than expected (p<0.001); --, less than expected (p<0.01); -, less than expected (p<0.05); NS, non-significant overlap.

**Supplementary Table S3. Genomic regions with correlated signal identified by Epicorr v1.20**

| *Region1* [a] | *Gene* | *Region2* (*Dido1*) [a] | *Score* [b] | *Description* [c] | *Chromatin states* [d] |
|---|---|---|---|---|---|
| 1:27,038,001-27,039,000 | *ARID1A* | 20:61,545,001-61,546,000 | 0.782 | SWI/SNF complex, chromatin remodelling | Enh, EnhG, Tx, TxWk |
| 2:60,772,001-60,773,000 | *BCL11A* | 20:61,545,001-61,546,000 | 0.8 | TF associated with the SWI/SNF complex | Tx, TxWk |
| 2:27,015,001-27,016,000 | *CENPA* | 20:61,545,001-61,546,000 | 0.704 | Centromere-associated protein network | Tx, TxWk |
| 4:104,089,001-104,090,000 | *CENPE* | 20:61,545,001-61,546,000 | 0.726 | Centromere-associated protein network | Tx, TxWk |
| 1:214,811,001-214,812,000 | *CENPF* | 20:61,545,001-61,546,000 | 0.809 | Centromere-associated protein network | Enh, Tx, TxWk |
| 1:214,814,001-214,815,000 | *CENPF* | 20:61,545,001-61,546,000 | 0.778 | Centromere-associated protein network | Tx, TxWk |
| 1:214,812,001-214,813,000 | *CENPF* | 20:61,545,001-61,546,000 | 0.742 | Centromere-associated protein network | Enh, Tx, TxWk |
| 1:214,805,001-214,806,000 | *CENPF* | 20:61,545,001-61,546,000 | 0.734 | Centromere-associated protein network | Enh, Tx, TxWk |
| 1:214,819,001-214,820,000 | *CENPF* | 20:61,545,001-61,546,000 | 0.729 | Centromere-associated protein network | Tx, TxWk |
| 2:234,759,001-234,760,000 | *HJURP* | 20:61,545,001-61,546,000 | 0.708 | H3 chaperone complex of CENP-A variant | Tx, TxWk |
| 2:234,747,001-234,748,000 | *HJURP* | 20:61,545,001-61,546,000 | 0.704 | H3 chaperone complex of CENP-A variant | EnhG, Tx, TxWk |
| 10:126,832,001-126,833,000 | *CTBP2* | 20:61,550,001-61,551,000 | 0.857 | Polycomb accessory protein (PRC2) | Enh, TxWk |
| 10:126,820,001-126,821,000 | *CTBP2* | 20:61,550,001-61,551,000 | 0.743 | Polycomb accessory protein (PRC2) | Enh, TxWk |

[a] Genomic regions, represented by chromosome number and start–end positions. [b] Epicorr v1.20 correlation score (>0.7). [c] Protein annotations from UniProtKB (www.uniprot.org). [d] Chromatin states overlapping between the two correlated regions. Chromatin state abbreviations: Enh, enhancers; EnhG, genic enhancers; Tx, strong transcription; TxWk, weak transcription.

**Supplementary Table S4. Cancer types, number of tumor and normal samples, and statistical significance of gene and protein expression differences**

| TCGA | Cancer type | Gene expression (TPM) | | | Protein expression (Z-score) | | |
|---|---|---|---|---|---|---|---|
| | | Tumour | Normal | Statistical significance | Tumour | Normal | Statistical significance |
| ACC | Adrenocortical carcinoma | 79 | 0 | n.a. | 0 | 0 | n.a. |
| AML | Acute myeloid leukaemia | 173 | 0 | n.a. | 0 | 0 | n.a. |
| BLCA | Bladder urothelial carcinoma | 408 | 19 | 2.5 x $10^{-01}$ | 0 | 0 | n.a. |
| BRCA | Breast invasive carcinoma | 1097 | 114 | 2 x $10^{-02}$ | | | |
| | Breast cancer | | | | 125 | 18 | **5.3 x $10^{-30}$** |
| CESC | Cervical squamous cell carcinoma | 305 | 3 | 2.9 x $10^{-01}$ | 0 | 0 | n.a. |
| CHOL | Cholangiocarcinoma | 36 | 9 | **3.6 x $10^{-11}$** | 0 | 0 | n.a. |
| | Clear cell renal cell carcinoma | | | | 110 | 84 | **9.8 x $10^{-11}$** |
| COAD | Colorectal adenocarcinoma | 286 | 41 | **9.9 x $10^{-13}$** | | | |
| | Colon cancer | | | | 97 | 100 | **2.7 x $10^{-32}$** |
| DLBC | Lymphoid Neoplasm Diffuse Large B-cell Lymphoma | 48 | 0 | n.a. | 0 | 0 | n.a. |
| ESCA | Oesophageal carcinoma | 184 | 11 | **6.8 x $10^{-03}$** | 0 | 0 | n.a. |
| | Gastric cancer | | | | 80? | 0 | n.a. |
| GBM | Glioblastoma multiforme | 156 | 5 | 4.2 x $10^{-01}$ | 99 | 10 | **3.6 x $10^{-18}$** |
| | Pediatric Brain Cancer | | | | 167? | 0 | n.a. |
| HNSC | **Head and neck squamous cell carcinoma** | 520 | 44 | **1.9 x $10^{-04}$** | 108 | 71 | **1.9 x $10^{-18}$** |
| | Hepatocellular carcinoma | | | | 165 | 165 | **3.1 x $10^{-57}$** |
| KICH | Kidney chromophobe carcinoma | 67 | 25 | **9.6 x $10^{-05}$** | 0 | 0 | n.a. |
| KIRC | Kidney renal clear cell carcinoma | 533 | 72 | 2.6 x $10^{-01}$ | 0 | 0 | n.a. |
| KIRP | Kidney renal papillary cell carcinoma | 290 | 32 | 9.3 x $10^{-01}$ | 0 | 0 | n.a. |
| LGG | Brain low-grade glioma | 513 | 0 | n.a. | 0 | 0 | n.a. |
| LIHC | Liver hepatocellular carcinoma | 371 | 50 | **1.6 x $10^{-12}$** | 0 | 0 | n.a. |
| LUAD | **Lung adenocarcinoma** | 515 | 59 | **1.2 x $10^{-04}$** | 111 | 111 | **1.3 x $10^{-28}$** |
| LUSC | Lung squamous cell carcinoma | 503 | 52 | 3.9 x $10^{-01}$ | 110 | 102 | **5.7 x $10^{-36}$** |
| MESO | Mesothelioma | 87 | 0 | n.a. | 0 | 0 | n.a. |
| OV | Ovarian serous cystadenocarcinoma | 305 | 0 | n.a. | | | |
| | Ovarian cancer | | | | 100 | 25 | **5.3 x $10^{-06}$** |
| PAAD | Pancreas adenocarcinoma | 178 | 4 | 3.9 x $10^{-01}$ | 137 | 74 | **1.7 x $10^{-09}$** |
| PCPG | Pheochromocytoma and paraganglioma | 179 | 3 | 1.9 x $10^{-01}$ | 0 | 0 | n.a. |
| PRAD | Prostate adenocarcinoma | 497 | 52 | **5.6 x $10^{-03}$** | 75? | 0 | n.a. |
| READ | Rectal adenocarcinoma | 166 | 10 | **1.7 x $10^{-03}$** | 0 | 0 | n.a. |
| SARC | Sarcoma | 260 | 2 | **< 1 x $10^{-12}$** | 0 | 0 | n.a. |
| SKCM | Skin cutaneous melanoma | 472 | 1 | n.a. | 0 | 0 | n.a. |
| STAD | Stomach adenocarcinoma | 415 | 34 | **2.1 x $10^{-07}$** | 0 | 0 | n.a. |
| TGCT | Testicular germ cell tumours | 150 | 0 | n.a. | 0 | 0 | n.a. |
| THCA | Thyroid carcinoma | 505 | 59 | **1.9 x $10^{-12}$** | 0 | 0 | n.a. |
| THYM | Thymoma | 120 | 2 | 1.6 x $10^{-02}$ | 0 | 0 | n.a. |
| UCEC | **Uterine corpus endometrial carcinoma** | 546 | 35 | **1.6 x $10^{-12}$** | 100 | 31 | 4.9 x $10^{-02}$ |
| UCS | Uterine carcinosarcoma | 57 | 0 | n.a. | 0 | 0 | n.a. |
| UVM | Uveal melanoma | 80 | 0 | n.a. | 0 | 0 | n.a. |

Statistical significance (P-values) for differences in gene (TCGA) and protein (CPTAC) expression levels between normal and tumor samples was calculated using Student's t-test via the UALCAN web portal (http://ualcan.path.uab.edu). Transcripts per million (TPM) values with significant differences (P<0.001) are highlighted in bold. Z-values represent standard deviations from the median across samples for the given cancer type. Log2 spectral count ratio values from CPTAC were normalized first within each sample profile, then across samples. Cancer type abbreviations follow those used by The Cancer Genome Atlas (TCGA; https://gdc.cancer.gov/resources-tcga-users/tcga-code-tables/tcga-study-abbreviations).

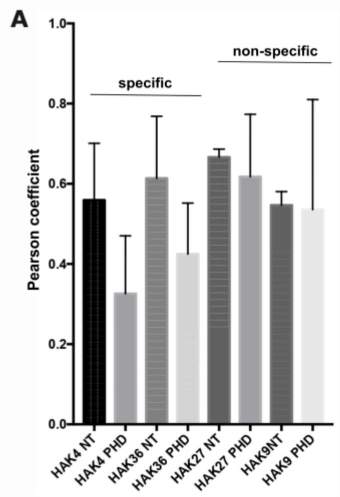


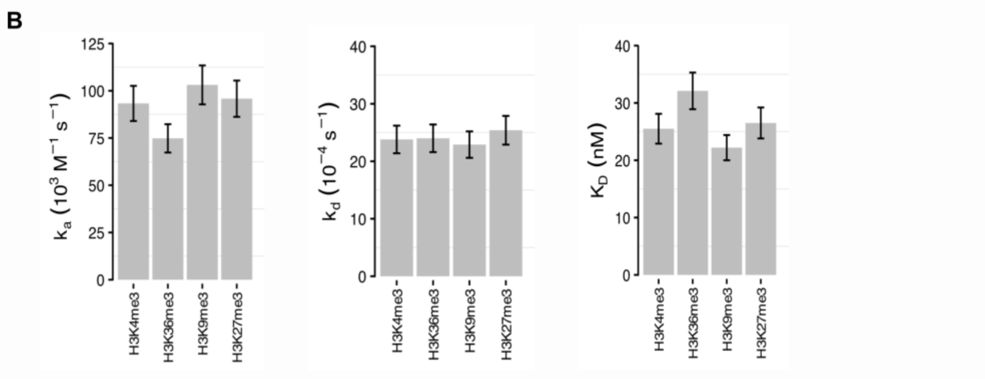


**C**

| | | Peptide | KD ($10^{-9}$M) | ka ($10^3$ $M^{-1}$ $s^{-1}$) | kd ($10^{-4}$ $s^{-1}$) | Method | Reference |
|---|---|---|---|---|---|---|---|
| DIDO3-PHD | H3K4 | ARTK(me3)QTARKST-Cys | 25.5 | 93.3 | 23.8 | SPRi | |
| DIDO3-PHD | H3K36 | ATGGVK(me3)KPHRY-Cys | 32.1 | 74.8 | 24.0 | SPRi | This study |
| DIDO3-PHD | H3K9 | KQTARK(me3)STGGK-Cys | 22.2 | 103.1 | 22.9 | SPRi | |
| DIDO3-PHD | H3K27 | TKAARK(me3)SAPAT-Cys | 26.5 | 95.8 | 25.4 | SPRi | |
| | | | KD ($10^{-6}$M) | | | | |
| BPTF-PHD | H3K4 | ARTK(me3)QTARKSTGGKAY | 3.0 | | | SPRi | DOI:10.1016/j.molcel.2007.10.023 |
| PYGO-PHD | H3K4 | ARTK(me3)QTARKSTGGKA | 2.5 | | | ITC | DOI:10.1016/j.molcel.2008.03.011 |
| TAF3-PHD | H3K4 | ARTK(me3)QTARKSTGG | 0.31 | | | ITF | DOI:10.1016/j.str.2008.04.015 |
| ATRX-PHD | H3K9 | ARTKQTARK(me3)STGGKA | 0.85 | | | SPRi | DOI:10.1038/nsmb.2062 |
| DPF3b-PHD | H3K14 | ARTKQTARKSTGGK(Ac)APRK | 1.2 | | | FPA | DOI:10.1038/nature09139 |
| RAG2-PHD | H3K4 | ARTK(me3)QTARKSTGGKY | 33.8 | | | FAA | DOI:10.1073/pnas.0709170104 |
| RAG2-PHD | H3R2/K4 | AR(me2)TK(me3)QTARKSTGGKY | 25.1 | | | FAA | DOI:10.1073/pnas.0709170104 |

**Supplementary Figure S1. DIDO3 co-localization and binding with the histone H3 tail**

(**A**) Pearson's coefficients for the co-localization of DIDO proteins with various histone H3 lysine trimethylation marks, calculated as detailed in Appendix 1, based on the degree of overlap between fluorescence signals in confocal microscopy images. (**B**) Surface plasmon resonance imaging (SPRi) analysis of the interaction between the DIDO3 PHD domain and histone-derived peptides. The isolated recombinant PHD domain was assayed for binding to immobilized H3K4me3, H3K9me3, H3K27me3, and H3K36me3 peptides. Error bars represent the standard error of the mean from three independent experiments. (**C**) Comparison of DIDO3 PHD domain affinity values with those of other PHD domains. Abbreviations: ITC, isothermal titration calorimetry; ITF, intrinsic tryptophan fluorescence binding experiments; FPA, fluorescence polarization assay; FAA, fluorescence anisotropy-based binding analysis.

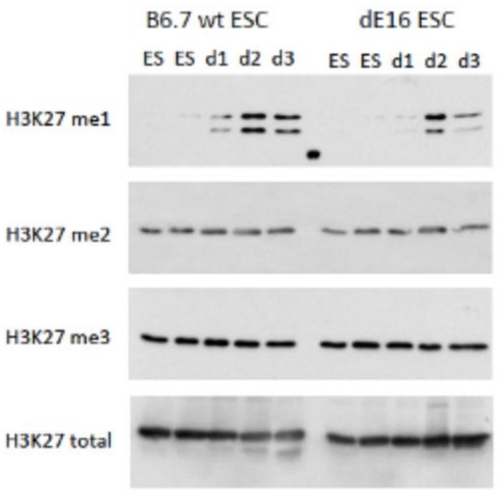


**Supplementary Figure S2. Methylation status of H3K27**

Histone lysates from wild-type (wt) and Dido3ΔE16 (dE16) ESCs and EBs (B6.7) at early differentiation (days 1–3) were probed with antibodies specific for mono-, di-, and trimethylated H3K27, and with an antibody against total H3K27.

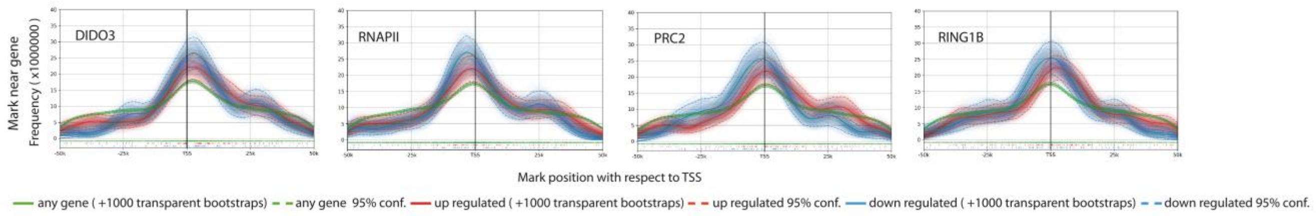


**Supplementary Figure S3. Distribution of DIDO3, RNAPII, PRC2 and PRC1 binding sites relative to the transcription start site of upregulated and downregulated genes**

The observed frequency of DIDO3, RNAPII, PRC2 and PRC1 binding sites around the transcription start site (TSS; from 50 kb upstream to 50 kb downstream) of upregulated and downregulated genes was compared to the frequency expected under a random distribution. The following datasets from the Gene Expression Omnibus (GEO) were used: GSE85029 (DIDO3 ChIP-seq), GSM2645517 (RNAPII ChIP-seq, largest subunit RPB1), GSE120393 (PRC2 ChIP-seq), GSM2645509 (PRC1 ChIP-seq, subunit RING1B), and GSE152346 (RNA-seq of DIDO3 and DIDO3ΔE16 in mouse embryonic stem cells).

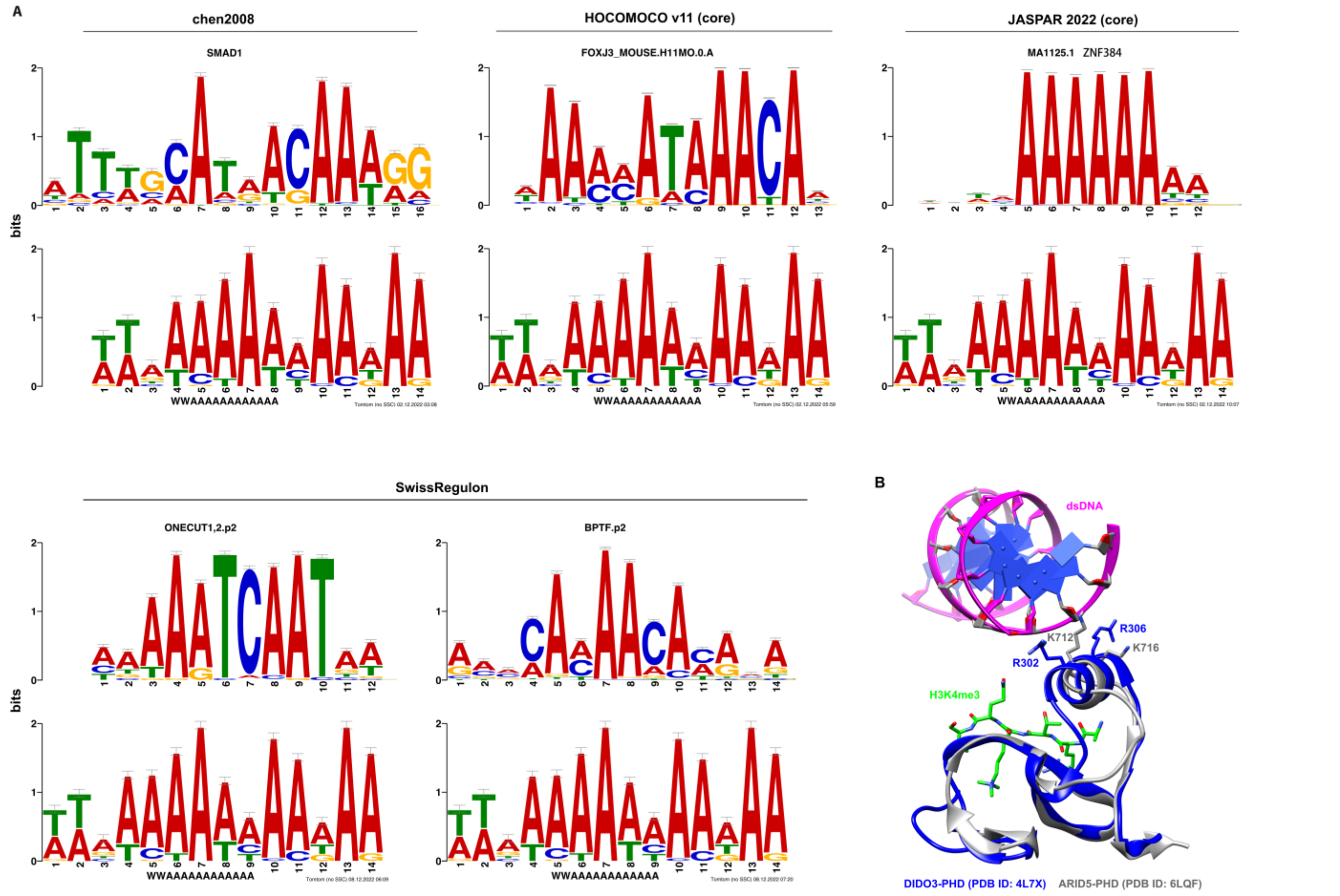


**Supplementary Figure S4. Overlap between DIDO3 binding sites and known transcription factor binding sites in the promoter regions**

(**A**) Sequence logo of the motif discovered by MEME-ChIP (bottom), matched against a known transcription factor (TF) DNA-binding motif (top). (**B**) Structural comparison of DIDO3-PHD in complex with an H3K4me3 peptide (PDB ID: 4L7X) and ARID5-PHD in ternary complex with an H3K4me3 peptide and AT-rich DNA (PDB ID: 6LQF).

**Supplementary File 1. Summary of the RNA-seq analysis in mouse embryonic stem cells (GSE152346) and mouse embryonic fibroblasts (NCBI Sequence Read Archive accession numbers SRP150516 and PRJNA476070)**

[tab-delimited]
Row names: row number
Columns:

- gene_symbol: Entrez gene symbol
- transcript_ID: UCSC transcript identifier
- exons: Annotated transcript exons
- dE16_log2FC: Log2 fold change in gene expression between DIDO3ΔE16 mutant and WT ESCs
- dE16_FDR: Adjusted p-value for dE16_log2FC
- dCT_log2FC: Log2 fold change in gene expression between DIDO3ΔCT+HA-DIDO3 mutant and WT ESCs
- dCT_FDR: Adjusted p-value for dCT_log2FC

**Supplementary File 2. ChIPseeker annotations for the DIDO3 ChIP-seq dataset**

[tab-delimited]
Row names: row number
Columns:

- seqnames: Chromosome name
- start: Start position of the peak in the genome
- end: End position of the peak in the DNA
- width: Length of the predicted binding peak
- info: Unique MACS2 identifier corresponding to a ChIP-seq peak from the DIDO3 dataset (GSE85029)
- annotation: Genomic region overlapping the ChIP-seq peak (i.e., Promoter, Exon, Intron, Downstream, Distal Intergenic, 3' UTR, 5' UTR)
- geneStrand: Strand direction of the gene nearest to a ChIP-seq peak (forward [1] or reverse [2])
- geneId: Entrez identifier for the gene nearest to a given ChIP-seq peak
- transcriptId: UCSC transcript identifier
- distanceToTSS: Distance from the ChIP-seq peak to the gene transcription start site (TSS)
- ENSEMBL: Ensembl gene identifier
- SYMBOL: Entrez gene symbol
- flank_txIds: UCSC transcript identifiers for the flanking genes
- flank_gene_EntrezIds: Entrez gene identifiers for the flanking genes
- flank_symbol: Entrez gene symbols for the flanking genes

**Supplementary File 3. Gene Ontology enrichment analysis for the genes within the correlated regions identified by Epicorr v1.20**

[tab-delimited]
Row names: row number
Sheet1 Columns:

- GO.ID: Gene Ontology identifier
- Term: Gene Ontology term description
- Annotated: Number of genes in the genome annotated with this GO term
- Significant: Number of genes in the input gene list annotated with this GO term
- Expected: Expected number of significant genes under a random distribution
- Fisher Test: P-value from Fisher's exact test for enrichment of this GO term

Sheet2 (Gene_list_1): List of genes with a correlation score > 0.7, identified by Epicorr v1.20
Sheet3 (Gene_list_2): List of genes with a correlation score between 0.6 and 0.7, identified by Epicorr v1.20

**Supplementary File 4. *Dido1* genomic alterations annotated in TCGA PanCancer Atlas studies**

[tab-delimited]
Row names: row number
Columns:

- Study: Name of the cancer study, as stored in cBioPortal
- Sample ID: Identification code of the sample

- Cancer Type: Type of cancer studied
- Protein Change: Protein consequence of the DNA change
- Annotation: Integrated resources for variant annotation. Prior knowledge about variants, including clinical actionability information, is provided from three sources: 1) OncoKB (www.oncokb.org), 2) CIViC (civicdb.org), and 3) My Cancer Genome (mycancergenome.org). Variants are also annotated as "hotspots" if the amino acid position was found to be a recurrent linear hotspot, as defined by the Cancer Hotspots method (cancerhotspots.org), or a three-dimensional hotspot, as defined by 3D Hotspots (3dhotspots.org)
- Functional Impact: Methods used to predict the functional impact of the variants
- Mutation Type: Type of change in DNA, or translational effect of the variant allele (e.g., missense, nonsense, frame shift deletion/insertion, in-frame deletion/insertion, splice region/site, fusion)
- Variant Type: Short code for the mutation type (i.e., SNP: single nucleotide variant; DEL: deletion; INS: insertion; NA: not annotated)
- Copy #: Type of copy number alteration (i.e., diploid, gain, shallowdel: shallow deletion, amp: amplification, deepdel: deep deletion)
- COSMIC: Number of occurrences of mutations at the same amino acid position in the COSMIC database
- Chromosome: Chromosome where the variant is located
- Start Pos: Start position of the variant in the genome
- End Pos: End position of the variant in the genome
- Ref: Wild-type nucleotide
- Var: Mutant nucleotide
- HGVSg: HGVS genomic nomenclature
- Allele Freq (T): Variant allele frequency in the tumor sample
- Allele Freq (N): Variant allele frequency in the normal sample
- # Mut in Sample: Number of mutations in the sample
